# Diversity and Function of the Eastern Oyster (*Crassostrea virginica*) Microbiome

**DOI:** 10.1101/2020.09.08.288811

**Authors:** Zachary T. Pimentel, Keith Dufault-Thompson, Kayla T. Russo, Abigail K. Scro, Roxanna M. Smolowitz, Marta Gomez-Chiarri, Ying Zhang

**Affiliations:** Department of Cell and Molecular Biology, University of Rhode Island, Kingston, RI, USA; Aquatic Diagnostic Laboratory, Roger Williams University, Bristol, RI, USA; Department of Fisheries, Animal and Veterinary Science, University of Rhode Island, Kingston, RI, USA

## Abstract

Marine invertebrate microbiomes play important roles in various host and ecological processes. However, a mechanistic understanding of host-microbe interactions is so far only available for a handful of model organisms. Here, an integrated taxonomic and functional analysis of the microbiome of the eastern oyster, *Crassostrea virginica*, was performed using 16S rRNA gene amplicon profiling, shotgun metagenomics, and genome-scale metabolic reconstruction. A relatively low number of amplicon sequence variants (ASVs) were observed in oyster tissues compared to water samples, while high variability was observed across individual oysters and among different tissue types. Targeted metagenomic sequencing of the gut microbiota led to further characterization of a dominant bacterial taxon, the class *Mollicutes*, which was captured by the reconstruction of a metagenome-assembled genome (MAG). Genome-scale metabolic reconstruction of the oyster *Mollicutes* MAG revealed a reduced set of metabolic functions and a high reliance on the uptake of host-derived nutrients. A chitin degradation and an arginine deiminase pathway were unique to the MAG as compared to other closely related *Mycoplasma* genomes, indicating a distinct mechanism of carbon and energy acquisition by the oyster-associated *Mollicutes*. A systematic reanalysis of public eastern oyster-derived microbiome data revealed the *Mollicutes* as a ubiquitous taxon among adult oysters despite their general absence in larvae and biodeposit samples, suggesting potential horizontal transmission via an unknown mechanism.

**IMPORTANCE:** Despite well-documented biological significance of invertebrate microbiomes, a detailed taxonomic and functional characterization is frequently missing from many non-model marine invertebrates. By using 16S rRNA gene-based community profiling, shotgun metagenomics, and genome-scale metabolic reconstruction, this study provides an integrated taxonomic and functional analysis of the microbiome of the eastern oyster, *Crassostrea virginica*. Community profiling revealed a surprisingly low richness, as compared to surrounding seawater, and high variability among different tissue types and individuals. Reconstruction of a *Mollicutes* MAG enabled the phylogenomic positioning and functional characterization of the oyster-associated *Mollicutes*. Comparative analysis of the adult oyster gut, biodeposits, and oyster larvae samples indicated the potentially ubiquitous associations of the *Mollicutes* taxon with adult oysters. To the best of our knowledge, this study represented the first metagenomics derived functional inference of the eastern oyster microbiome. An integrated analytical procedure was developed for the functional characterization of microbiomes in other non-model host species.

## INTRODUCTION

Oysters are filter feeding bivalve molluscs with great ecological and economic significance. They are considered ecosystem engineers due to their ability to form reefs that serve a variety of beneficial functions, including protecting shorelines from storm-related damage and providing habitat for other marine organisms [1, 2]. Other important environmental functions provided by both wild and farmed oysters include biological remediation, sequestering of carbon through calcium carbonate shell productions, and alteration of biogeochemical cycles (such as the promotion of microbially mediated denitrification process) through their filter feeding lifestyle that involves deposition of feces and pseudofeces into the sediments [3, 4]. For these reasons, efforts have been made to restore oyster populations around the United States. In addition, oysters represent a significant and growing portion of the aquaculture industry. In 2018, *Crassostrea* spp. of oysters represented almost one-third of the major species produced in world aquaculture [5].

Like other invertebrates, oysters are known to harbor a diverse range of microorganisms. Culture-based studies of oyster-associated microbes have revealed the presence of *Proteobacteria* (e.g. *Vibrio*, *Pseudomonas*, *Alteromonas*), *Actinobacteria* (e.g. *Micrococcus*), and *Bacteroidetes* (e.g. *Flavobacterium*) [6]. Some readily culturable members of the oyster microbiome, such as strains of the *Vibrio* genus, have been closely studied to profile their abundance [7], pathogenic potential [8], evolution and diversity [9], and inhibition of growth by probiotics [10, 11]. Other well known, cultured microbes from oysters include the protozoan pathogen *Perkinsus marinus*, a causative agent of Dermo disease, which is one of the major diseases of adult eastern oysters [12]. Prior research has identified epizootiological factors involved with the prevalence and immunological consequences of the *P. marinus* infection [13, 14]. However, little difference has been observed in the diversity and composition of the oyster-associated bacterial communities during *P. marinus* infection [15, 16].

Culture-independent approaches, such as the profiling of amplicon libraries, lead to the detection of other previously uncultured taxa in the oyster microbiome, such as the *Chloroflexi*, *Firmicutes*, *Fusobacteria*, *Planctomycetes*, *Spirochaetes*, *Tenericutes*, and *Verrucomicrobia* [17–19]. A distinct microbiome is found in multiple oyster tissues (e.g. hemolymph, gill, mantle, and gut) when compared with microbial communities of the surrounding seawater, suggesting potential host selections that lead to the enrichment of specific groups [20]. Multiple factors, including changes in environmental conditions [15,17,21,22], diet [23], infection [24–26], and the use of probiotics [27], have been shown to influence the composition of oyster microbiomes during certain life stages and among different tissue types. All these studies set the stage for further investigating the taxonomic composition and functional potentials of the oyster microbiomes across different tissues.

In the absence of microbial isolates, shotgun metagenomics serves as a useful tool for gaining functional insights into uncultured members of a microbiome. Prior applications of metagenomics in marine invertebrates have revealed remarkable bacterial functions, including chemical defense mediated by secondary metabolites produced by the sponge microbiome [28] and the metabolic interactions between chemosynthetic symbionts and their hosts in deep-sea hydrothermal vents [29]. One potential challenge in the application of metagenomics to host tissues is the high abundance of host DNA that masks the signals from the tissue-associated microbiome [30]. This issue may not be easily resolved by targeting different sample types (e.g. using biodeposit samples to represent the gut microbiota), as a clear distinction is found between the fecal microbiome and the gut microbiome of filter feeding bivalves, e.g. the blue mussel, *Mytilus edulis* [31]. Thus, a successful application of metagenomics in oyster microbiome studies requires the development of customized protocols for the enrichment of microbial DNA from the oyster host tissues.

Here, an integrated microbiome analysis of the eastern oyster, *Crassostrea virginica*, was performed by combining 16S rRNA gene-based community profiling, shotgun metagenomics, and genome-scale metabolic reconstruction. Amplicon libraries of six distinct oyster tissue types were analyzed to compare the diversity and distribution of microbiomes among individual oysters. Metagenomic-based identification of oyster gut-associated bacteria was enabled with a specialized protocol that enriched microbes from host tissues. Functional characterization was performed through the application of a genome-scale metabolic reconstruction on a metagenome-assembled genome (MAG).

## RESULTS

### Sample Descriptions

Tissue samples were dissected from 60 farmed oysters collected from Ninigret Pond, Rhode Island, United States of America. In order to evaluate potential relationships between bacterial profiles and oyster health status, individual oysters were tested for potential infection by the most common causative agent of oyster disease in the region, the parasite *Perkinsus marinus*, using a real-time PCR assay (**Materials and Methods**). Overall, 14 of the 60 oysters were infected with *P. marinus* with a range of infection severities measured by the Mackin Index, an indicator of infection intensity based on cycle threshold values from the real-time PCR assay [32]. Subsets of infected and uninfected oysters were selected for further molecular profiling of the microbiome. Microbial community profiling was performed for 19 oysters on the gut, mantle, gill, and inner shell swab samples and for a subset of 9 oysters on the pallial fluid and hemolymph samples. As a comparison to the oyster tissues, samples were taken from the surrounding water column to profile the free-living (0.2-5 µm) and particle-associated (5-153 µm) microbial communities. Additionally, gut samples from 12 oysters were prepared for barcoded metagenomic sequencing, and pooled gut samples from another 10 oysters were prepared with an enrichment procedure for microbiome-enriched metagenomic sequencing (**Materials and Methods**). A summary of all oyster samples, the *P. marinus* Mackin Indices, and sequenced tissue types was provided in **Supplemental Table S1**.

### Community Diversity of the Oyster Microbiome

The community richness, as measured by the Amplicon Sequence Variant (ASV) counts, was determined for samples of each tissue type (**Figure 1A**). The median number of ASVs per sample among the six tissue types ranged from 39 to 94, with the lowest counts in the gut and mantle and the highest counts in the inner shell swab (**Supplemental Table S2**). All oyster tissues had relatively lower richness compared to the surrounding seawater, as evidenced by the sample-based rarefaction curves (**Supplemental Figure S1**). According to the rarefaction curves, the oyster tissue samples converged well before a sampling depth of 5,000 ASVs, with the majority of samples containing less than 200 distinct ASVs. In contrast, taxonomic profiling of the water samples barely converged with a sampling depth of over 20,000 ASVs, with 296 and 752 ASVs, respectively, identified from the free-living and particle-associated microbial fractions.

**Figure 1.**
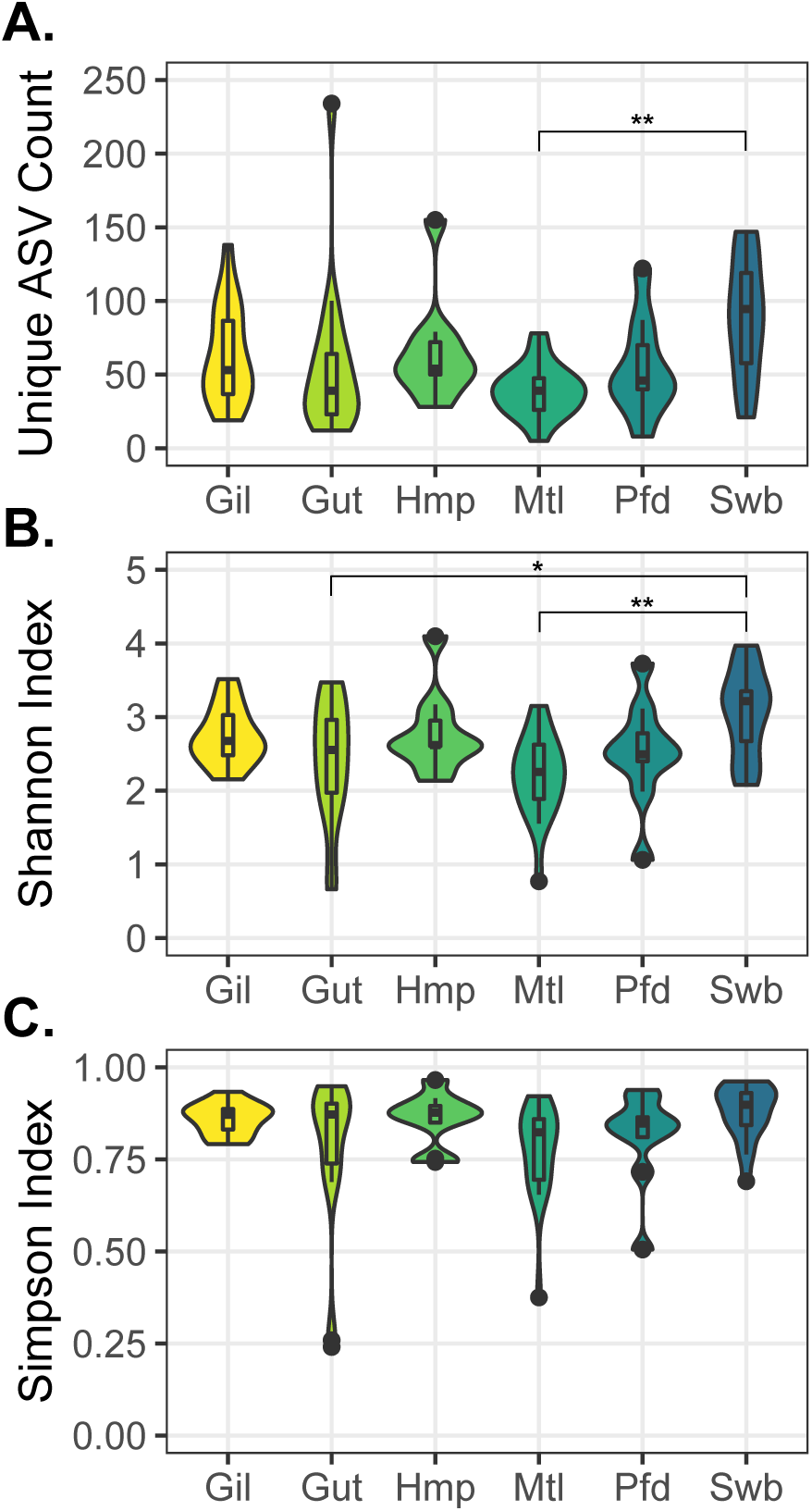
Diversity of eastern oyster microbiome samples as measured by (A) richness, based on the unique ASV count in each sample, (B) Shannon Index, and (C) Simpson Index. Samples were grouped by different tissue types, with acronyms as follows: Gil (Gill), Gut (Gut), Hmp (Hemolymph), Mtl (Mantle), Pfd (Pallial Fluid), and Swb (Inner Shell Swab). Pairwise statistical significance was assessed with Tukey’s Honest Significant Differences test: * p-value < 0.05, ** p-value < 0.01.

Despite the low number of ASVs across individual oyster tissue samples, relatively high variability was observed both among different tissue types and across different individuals. When comparing a tissue type across different individuals, only a small fraction of all observed ASVs were conserved (present in >80% of samples within a tissue type), with the shell swab representing the lowest percentage (1.3%) and the hemolymph representing the highest percentage (5.8%) of conserved ASVs (**Supplemental Table S2**). Similarly, when comparing the pooled ASVs from different oyster tissues, a total of 1,640 unique ASVs were identified among all the oyster samples, but only 50 ASVs were conserved among the six tissue types analyzed (**Supplemental Figure S2**). The inner shell swab samples were significantly more diverse when compared to the mantle samples based on richness (unique ASV counts) and Shannon index using the Tukey Honest Significant Differences test (TukeyHSD, p-value < 0.05) (**Figure 1A, 1B**). Similarly, a higher Shannon index was observed in the inner shell swab compared to the gut samples (**Figure 1B**). However, no statistical significance was detected among other tissue types. In addition, none of the tissue pairs showed any statistical difference based on the Simpson index (**Figure 1C**).

PERMANOVA analysis of the Bray-Curtis, Jaccard, Weighted UniFrac, and Unweighted UniFrac distances were performed to explore similarities and differences among samples of different tissue types (**Supplemental Figure S3**). Analysis of all four distance matrices suggested that the gut samples were statistically different from the rest of the profiled tissues (p-value < 0.05). Similarly, significant differences were observed between mantle samples and all other tissue types based on Bray-Curtis, Jaccard, and Weighted UniFrac distances. However, the comparisons between gill, pallial fluid, hemolymph, and inner shell swab samples resulted in different estimations of the statistical significance depending on the distance matrix used. While the gill samples were different from all other samples based on the Jaccard or Unweighted Unifrac distances, the differences between gill and pallial fluid samples were not supported when Bray-Curtis or Weighted UniFrac distances were examined.

### Taxonomic Identification of the Oyster Microbiome

Taxonomic assignment of the ASVs obtained from this study revealed several major bacterial classes that had a mean relative abundance of greater than 10% within at least one tissue type. These included *Gammaproteobacteria*, *Mollicutes*, *Fusobacteriia*, *Spirochaetia*, and *Chlamydiae*. Further, the *Gammaproteobacteria* was largely represented by the genus *Vibrio* in most tissue types (**Figure 2A,B**). The relative abundance of these major taxa was shown to vary among different tissue types based on the Tukey HSD test (p-value < 0.05). The genus *Vibrio* had a significantly lower relative abundance in gut samples as compared to other tissues (**Figure 2B**). In contrast, *Spirochaetia* had statistically higher presence in the mantle samples as compared to all other tissue types (**Figure 2D**). Meanwhile, the *Mollicutes* had statistically higher relative abundance in gut samples as compared to gill, mantle, and pallial fluid samples, but its higher mean abundance was not statistically significant when compared to the inner shell swab or hemolymph samples (**Figure 2E**). Another major taxon, *Fusobacteriia*, was relatively equally represented among all tissue types, with the mean relative abundances ranging from 11% in the mantle to 21% in the pallial fluid (**Figure 2C**). Finally, *Chlamydiae* had a higher mean relative abundance in the gut samples as compared to other tissue samples. However, this higher mean relative abundance was largely attributed to four gut samples in which the relative abundance of *Chlamydiae* ranged from 38% to 86% (**Figure 2F**). Meanwhile, the *Chlamydiae* were absent in over one-third of the gut samples.

**Figure 2.**
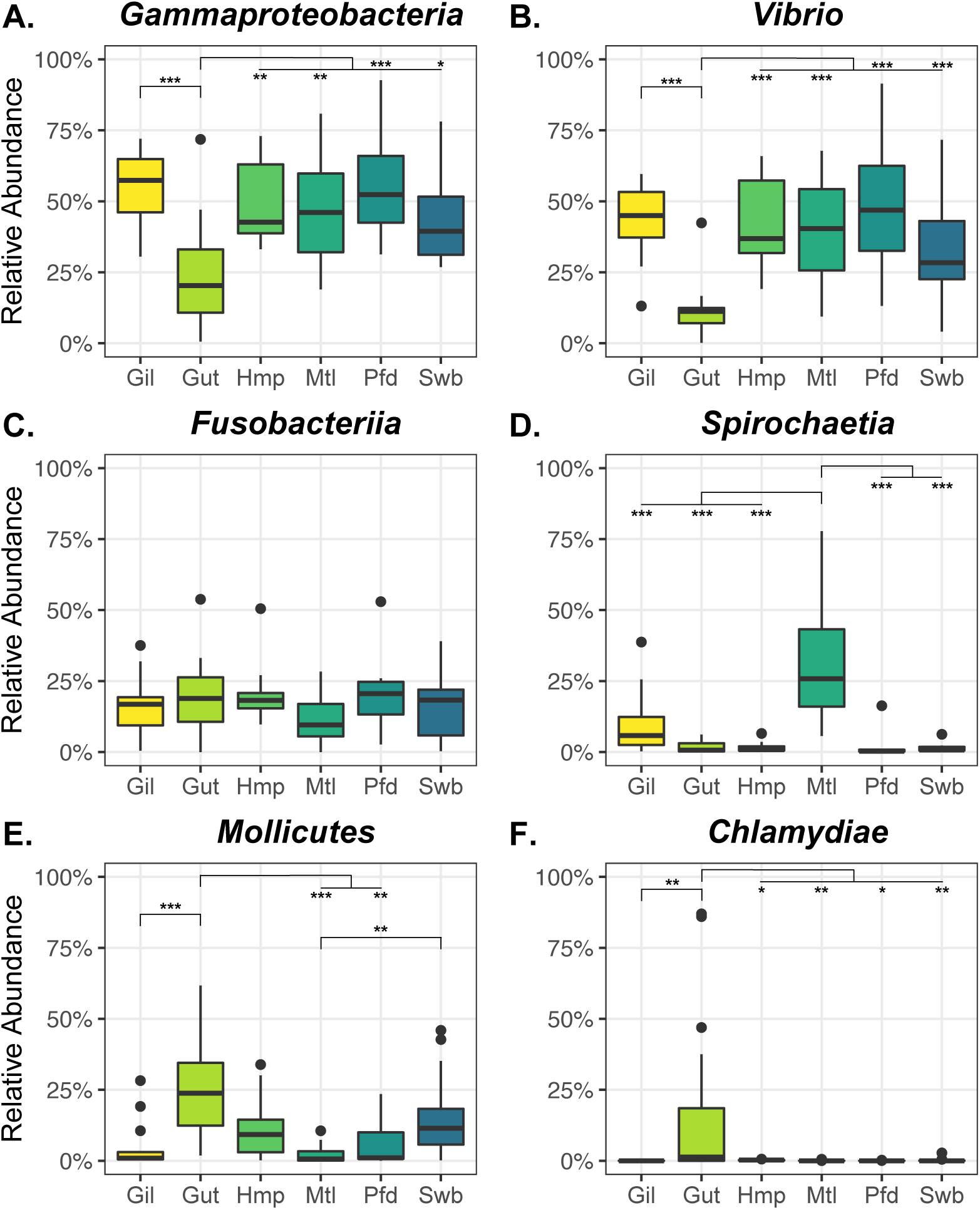
Boxplot showing the relative abundance of major taxa in individual samples grouped by tissue type: *Gammaproteobacteria* (A), *Vibrio* (B), *Fusobacteriia* (C), *Spirochaetia* (D), *Mollicutes* (E), and *Chlamydiae* (F). Each tissue type was given a three letter code as follows: Gil (Gill), Gut (Gut), Hmp (Hemolymph), Mtl (Mantle), Pfd (Pallial Fluid), and Swb (Inner Shell Swab). Pairwise statistical significance was assessed with Tukey’s Honest Significant Differences test: * p-value < 0.05, ** p-value < 0.01, *** p-value < 0.001.

Compared to their presence in the oyster tissue samples, most of the major taxa had low relative abundances in the surrounding water column. For example, the relative abundance of *Mollicutes*, *Chlamydiae*, *Fusobacteriia*, and *Spirochaetia* were below 0.7% in both the free-living fraction and the particle-associated fraction of the seawater (**Supplemental Figure S4**), which were orders of magnitude lower than the mean abundance of these taxa in some oyster tissues (**Figure 2**). The *Gammaproteobacteria* had a higher relative abundance than the other taxa, with 11% in the free-living fraction and 13% in the particle-associated fraction of the seawater. However, the *Vibrio* was not a dominant *Gammaproteobacteria* in the seawater. Compared to a mean *Vibrio* abundance of above 25% across most oyster tissues (except for the gut), the *Vibrio* in seawater had a relative abundance of 0.1% in the free-living fraction and 0.3% in the particle-associated fraction (**Figure 2** and **Supplemental Figure S4**).

### Metagenomic Sequencing of the Oyster Microbiome

Metagenomic sequencing was applied to the oyster gut samples in order to characterize the functional potential of the microbiome. Besides the barcoded sequencing of individual oyster samples, a procedure was adapted to enrich microbial signals and minimize genomic materials from the host (**Materials and Methods**). Overall, two rounds of metagenomic sequencing were performed, resulting in the collection of 13 metagenomes with a total of over 2.5 billion raw reads. Around 80% and 0.4% of the quality filtered reads were assigned to the oyster host and *P. marinus*, respectively. Besides these, a total of 394,986,464 reads were used for the assembly and binning of MAGs across the 13 metagenomes.

A co-assembly of these reads resulted in 612,574 unique contigs. At least one complete 16S rRNA gene was detected for each of the major taxa, ranging from 1,396 to 1,562 base pairs in length. Six of the full-length 16S rRNA genes had identical sequences (with 100% coverage) to the ASVs (from the V4 region) from amplicon sequencing, with taxonomy mappings to the *Mollicutes* (*Mollicutes-1* and *Mollicutes-2*), *Chlamydiae*, *Spirochaetia* (*Spirochaetia-1* and *Spirochaetia-2*), and *Bacteroidia* (**Supplemental Table S3**). The relative abundance of these sequences was probed across different oyster tissues and between *P. marinus* infected and uninfected samples based on mappings to the ASV abundances (**Supplemental Figure S5**). The highest abundance was observed with ASVs mapped to the *Mollicutes-2* and *Spirochaetia-1*, with both presenting in different tissue types. The mean relative abundance of *Mollicutes-2* was around 10% in several tissue types and was significantly higher in the uninfected than the *P. marinus* infected oyster gill and pallial fluid samples. The mean relative abundance of *Spirochaetia-1* was around 20% in the mantle samples, while no significant differences were observed in its abundance between infected and uninfected oysters. Two sequences, *Mollicutes-1* and *Chlamydiae*, occurred mainly in the gut tissue, where the average relative abundance was around 5%. Both sequences had a visibly higher average relative abundance in uninfected gut samples as compared to the *P. marinus* infected tissues, but this difference was not statistically significant. Lastly, the *Spirochaetia-2* and *Bacteroidia* were present in only a few oyster tissues and had a low abundance throughout all samples.

Binning of the co-assembly produced two MAGs with completeness greater than 80% and contamination less than 2%. Each MAG contained a full-length 16S rRNA gene, with the first from the class *Mollicutes* (*Mollicutes-1*) and the second from *Chlamydiae*. The *Mollicutes* MAG included 47 contigs with a total length of 0.62 Mb and a GC content of 28.7%. The estimated completeness and contamination was 97.4% and 1.8%, respectively. Similarly, the *Chlamydiae* MAG included 57 contigs with a total length of 1.08 Mb and a GC content of 41.0%, and it had a completeness of 86.2% and contamination of 0.4%. Besides the *Mollicutes* and *Chlamydiae* MAGs, a partial *Spirochetes* MAG was reconstructed, with a completeness of 43.2%, a contamination of 0.7%, and a full-length 16S rRNA gene (*Spirochaetia-1*) included in the MAG.

Phylogenomic reconstructions were performed based on conserved single-copy genes (CSCGs) identified between the two near-complete oyster MAGs and corresponding reference genomes in the *Mollicutes* and *Chlamydiae* taxa (**Materials and Methods**). In total, 34 CSCGs and 179 CSCGs, respectively, were used to build the *Mollicutes* and *Chlamydiae* phylogenies. Based on the phylogenomic reconstruction, the oyster *Mollicutes* MAG formed a basal branch to a group of marine *Mycoplasma*, with the nearest neighboring branches including *Mycoplasma todarodis* (isolated from a squid), *Mycoplasma marinum* (isolated from an octopus), and *Mycoplasmatales* bacterium DT_67 and DT_68, two MAGs obtained from deep-sea sinking particles (hereafter referred to as DT_67 and DT_68) [33] (**Figure 3A**). Meanwhile, the Chlamydiae MAG was found within the order *Parachlamydiales* most closely related to *Simkania negevensis*, an obligate intracellular bacteria with a broad host range from amoeba to animals [34, 35] (**Supplemental Figure S6**).

**Figure 3.**
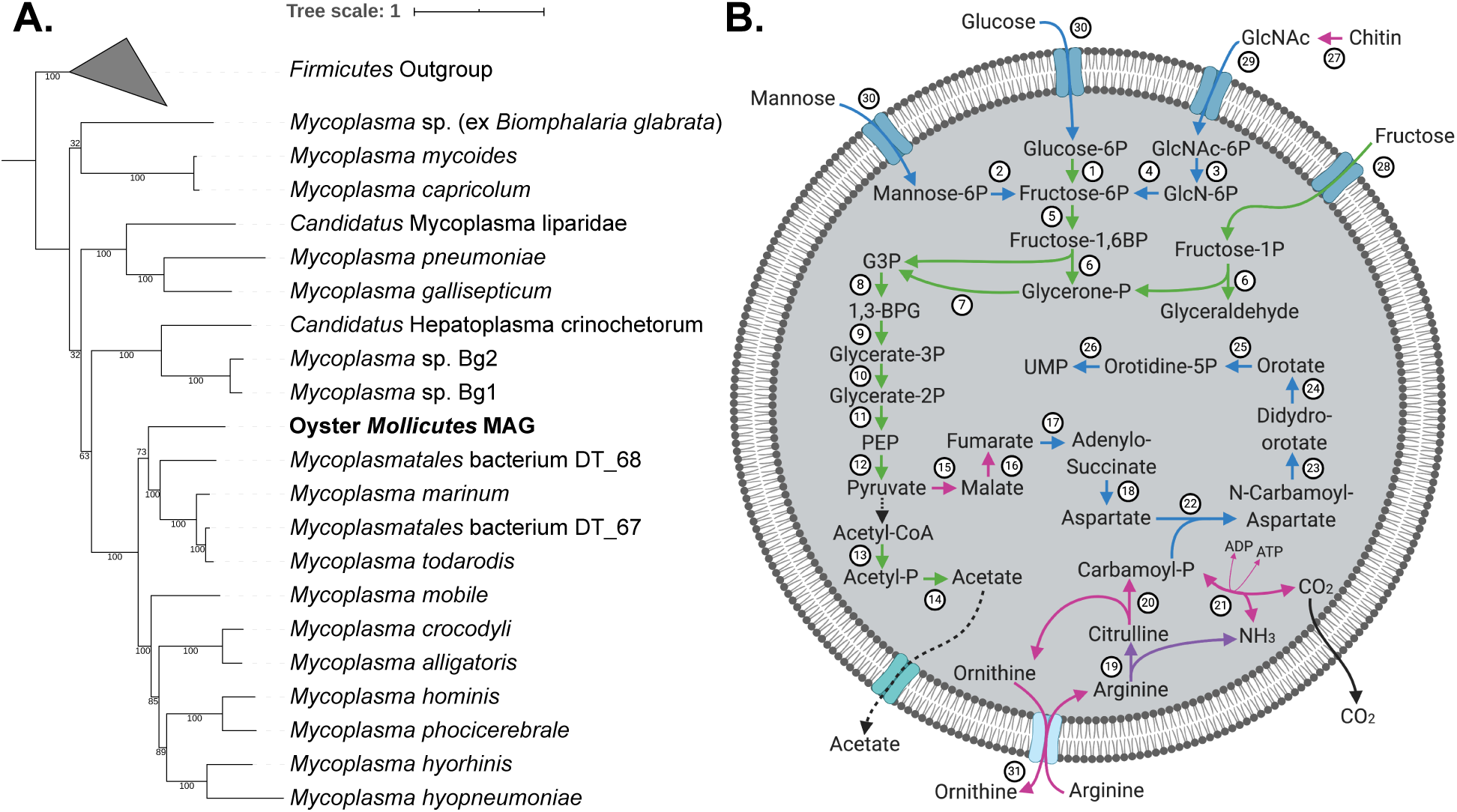
Phylogenomic and functional analysis of the oyster *Mollicutes* metagenome-assembled genome (MAG) identified from this study. **(A)** Maximum likelihood phylogenomic reconstruction based on conserved single-copy genes. Members of the *Firmicutes* were selected as outgroups in the phylogeny (**Supplemental Table S4**). Support values were based on 100 iterations of bootstrapping. (**B**) Visualization of the central metabolic pathways reconstructed from the oyster *Mollicutes* MAG (created with BioRender). Metabolites are connected with directed edges indicating biochemical conversions or transport and diffusion processes. The edges were coded by circled numbers, with further details represented in **Supplemental Table S6.** Additional coding was provided by the edge colors to indicate conservations between the oyster *Mollicutes* MAG and reference genomes, including the marine host-associated *Mycoplasma marinum* and *Mycoplasma todarodis*, and the freshwater host-associated *Mycoplasma mobile*. Green–conserved in all four genomes; blue–conserved between the *Mollicutes* MAG and the marine *Mycoplasma* (*M. marinum* and *M. todarodis*) but absent in *M. mobile*; purple–conserved between the *Mollicutes* MAG and the freshwater *M. mobile*, but absent from the marine *Mycoplasma*; magenta–unique functions in the *Mollicutes* MAG. The conversion from pyruvate to acetyl-CoA was marked as black because the function of a pyruvate dehydrogenase was identified outside of the MAG from unbinned contigs that had top BLAST hits to members of the *Mycoplasma*.

To determine the similarity of the *Mollicutes* and *Chlamydiae* MAGs to genomes within their corresponding taxa, average nucleotide identity (ANI) and average amino acid identity (AAI) were calculated between the MAGs and all other species represented in their corresponding CSCG phylogenies (**Supplemental Table S4, S5**). For the *Mollicutes* MAG, the highest pairwise ANI values were observed with *M. marinum* (66.8%), DT_67 (66.7%), DT_68 (66.4%), and *M. todarodis* (66.0%), all representing strains from marine sources, as well as *Mycoplasma mobile* (66.1%), a freshwater fish pathogen. Similarly, some of the highest pairwise AAI values were observed with *M. marinum* (51.9%) and *M. todarodis* (50.3%) with orthologs identified from 57.4% and 53.3%, respectively, of the protein-coding genes in the oyster *Mollicutes* MAG. While DT_67 and DT_68 had slightly higher AAI values of 54.4% and 53.6%, only a limited number of orthologs were identified, covering 22.5% and 22.2% of the protein-coding genes in the *Mollicutes* MAG (**Supplemental Table S4**). The oyster *Chlamydiae* MAG had an ANI between 62.8% and 64.8% to all reference genomes of the *Chlamydiae* phylum. The highest AAI value (47.9%) was observed with its nearest neighbor in the phylogenomic reconstruction, *Simkania negevensis*, with ortholog identifications for 53.6% of the protein-coding genes in the *Chlamydiae* MAG (**Supplemental Table S5)**. Overall, both the *Mollicutes* and *Chlamydiae* MAGs had lower ANI and AAI values to the reference genomes than expected for strains of the same species (both generally ≥ 95%) [36, 37].

### Novel Functional Potentials of the Oyster *Mollicutes* MAG

A genome-scale metabolic reconstruction was performed to illustrate the metabolic capacity encoded by the oyster *Mollicutes* MAG (**Figure 3B**). Overall, the *Mollicutes* MAG encoded a highly reduced metabolism with few carbon utilization pathways and limited biosynthetic capability. Transport via the PTS system was predicted for glucose, fructose, mannose, and N-acetylglucosamine (GlcNAc). A complete glycolysis pathway and all subunits of ATP synthase were present, while genes of the tricarboxylic acid (TCA) cycle were largely missing. The ability to convert pyruvate to fumarate was predicted based on the presence of a malate dehydrogenase and a fumarate hydratase. The conversion of acetyl-CoA to acetate was inferred by the identification of a phosphate acetyltransferase and an acetate kinase. An arginine deiminase (ADI) pathway was identified, potentially contributing to the production of ATP via the exchange of arginine and ornithine [38] and enabling the production of precursors for a *de novo* pyrimidine biosynthesis pathway (**Supplemental Table S6)**. The *Mollicutes* MAG also encoded a putative chitinase with the catalytic domain homologous to an experimentally verified chitinase in *Pyrococcus furiosus* (PDB: 2DSK) and a chitin-binding domain at the C-terminus. It was noted that a mechanism of converting pyruvate to acetyl-CoA was missing in the MAG. However, putative subunits of a pyruvate dehydrogenase complex were identified from other contigs in the metagenomic co-assembly, with top BLAST hits to members of the *Mycoplasma* genus. Therefore, the potential conversion of pyruvate to acetyl-CoA was speculated in the metabolic reconstruction (**Figure 3B**).

A comparative genomic analysis was performed between the oyster *Mollicutes* MAG and genomes from closely related *Mycoplasma* isolates, including two marine species, *M. marinum* and *M. todarodis*, as well as one freshwater species, *M. mobile*. Genes in the oyster *Mollicutes* MAG can be largely classified into four categories based on their conservation with other species from the comparative genomic analysis (**Supplemental Figure S7**). Over 40% of the genes in the *Mollicutes* MAG were conserved among all reference genomes, encoding functions in central metabolism (e.g. ATP synthase, amino acid metabolism, nucleotide metabolism, *etc*.) and information storage and processing (e.g. replication, translation, transcription). Around 10% of genes were conserved between the *Mollicutes* MAG and the two marine isolates but were missing from *M. mobile*. These included genes belonging to a *de novo* pyrimidine biosynthesis pathway, multiple carbohydrate utilization genes, and an alpha-amylase domain-containing protein. Only 18 genes (less than 3%) were uniquely conserved between the *Mollicutes* MAG and *M. mobile*, including an arginine deiminase and genes associated with DNA repair and the repair of enzymes under oxidative stress. Finally, around 36% of the genes were unique to the *Mollicutes* MAG. These included genes involved in pyruvate metabolism (e.g. malate dehydrogenase and fumarate hydratase) and the ADI pathway (e.g. carbamate kinase, ornithine carbamoyltransferase, and the arginine/ornithine antiporter). Overall, the oyster *Mollicutes* MAG maintained conserved functions with known *Mycoplasma* species while carrying unique metabolic capability in its genome.

### Presence and Distribution of *Mollicutes* across Eastern Oyster Microbiome Studies

Following functional inferences obtained from the metabolic reconstruction of the oyster *Mollicutes* MAG, emphasis was placed on the composition and diversity of *Mollicutes* across different oyster microbiomes obtained from this and other studies. A total of three additional datasets were identified from public databases, including the community profiling of gut samples from the adult eastern oyster [39], the profiling of adult eastern oyster biodeposits [40], and the profiling of eastern oyster larvae homogenate [27]. Raw data from these prior studies were downloaded and reanalyzed with a standardized pipeline to minimize potential inconsistencies caused by variation in computational procedures (**Materials and Methods**). According to the comparative analysis, a consistently high relative abundance of *Mollicutes* was found in adult oyster gut samples, but this taxon was largely absent from larvae homogenate and adult biodeposits. This is in contrast to other major taxa examined. For example, the *Gammaproteobacteria*, dominated by the *Vibrio* genus, had significantly higher relative abundance in larvae homogenate compared to the adult gut or biodeposits (**Supplemental Figure S8**).

To further elucidate the taxonomic distribution of the *Mollicutes* ASVs recovered from oyster tissues and the surrounding seawater, a phylogenetic analysis was performed by placing the *Mollicutes* ASVs from this study and a previous study of the adult eastern oyster gut microbiome [39] into a reference tree of 16S rRNA gene sequences from the SILVA database (**Materials and Methods**). Branches in the tree were collapsed with labels indicating the environmental source of the SILVA references. A bubble plot was included on the right of the phylogeny to indicate the number of unique *Mollicutes* ASVs identified from different tissue or water samples (**Figure 4**).

**Figure 4.**
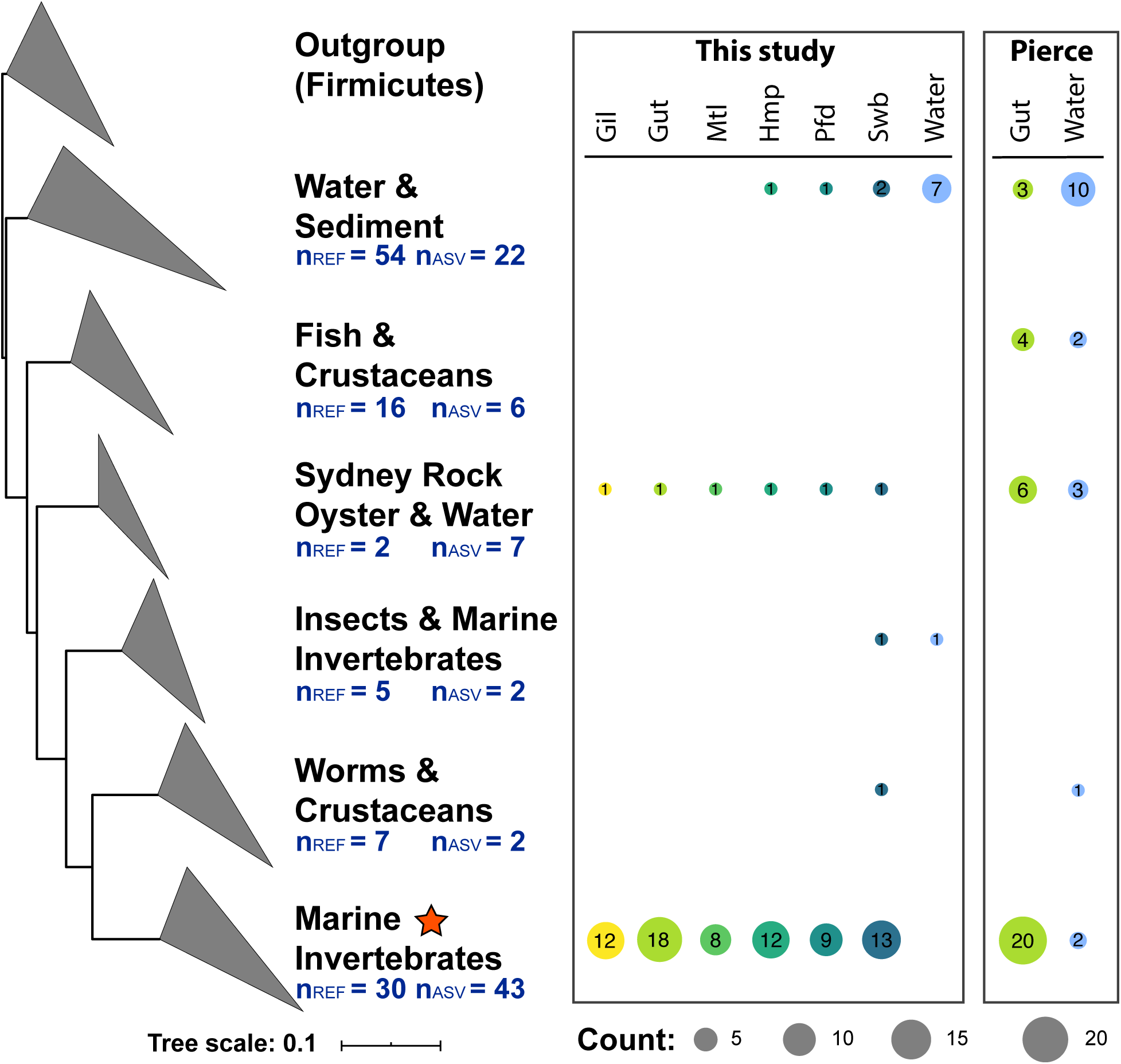
Phylogenetic positioning of oyster-associated *Mollicutes* ASVs. The branches were collapsed to summarize the environments represented by reference 16S rRNA genes in the SILVA database. The number of SILVA reference sequences (n_REF_) and unique ASV sequences (n_ASV_) were marked in each collapsed clade. Corresponding counts of unique ASVs in the different sample types were shown as a bubble plot for each clade on the right of the phylogeny, with filled circles sized according to the ASV counts and colored based on sample types. Data from this study and a prior study [39] was included in the analysis. Sample types included seawater the following oyster tissues: Gil (Gill), Gut (Gut), Hmp (Hemolymph), Mtl (Mantle), Pfd (Pallial Fluid), and Swb (Inner Shell Swab). A given ASV may appear in multiple sample types, and thus, the sum of values in each row of the bubble plot is not necessarily the number of ASVs (n_ASV_) in the collapsed branch. The clade labeled as Marine Invertebrates is marked with an orange star to indicate the primary clade of oyster-associated *Mollicutes*, which includes the two metagenome-assembled full-length 16S rRNA genes, *Mollicutes-1* and *Mollicutes-2*, and their corresponding ASVs (**Supplemental Table S3**).

The *Mollicutes* ASVs identified from oyster tissues had a largely distinct phylogenetic profile from the surrounding water samples. The tissue-associated ASVs consistently grouped around a clade dominated by references from diverse marine invertebrates, such as crabs, octopus, squid, and the Sydney rock oyster (*Saccostrea glomerata*) (marked with a star in **Figure 4**). In contrast, the water-associated ASVs grouped around a distant clade with references from water and sediments. Interestingly, a small number of ASVs from the oyster hemolymph, pallial fluid, and inner shell swab samples grouped within the clade of the water and sediment-associated *Mollicutes*, and the inner shell swab also included additional ASVs related to references in insects and marine invertebrates, such as coral, worms, and crustaceans. An additional clade, represented by only two references from the Sydney rock oyster and water, contained ASVs that are present in all the examined tissue types and some water samples. The phylogenetic profiling of *Mollicutes* ASVs in oyster tissues is well conserved between this study and the study by Pierce and Ward [39], where minor overlaps were found between ASVs from the oyster gut and the water samples. A detailed mapping of the Mollicutes references and ASVs to the different clades was shown in **Supplemental Table S7**.

## DISCUSSION

The microbiome of marine invertebrates has potential significance in mediating biological and ecological functions of host organisms [41, 42]. A comprehensive understanding of host-microbiome interactions relies on the taxonomic and functional characterization of the microbiome across different tissue types and individuals. Despite potential encounters with the highly diverse microbial communities in the surrounding water column through their filter feeding lifestyle, eastern oysters have been shown to generally contain low microbial diversity compared to the surrounding water column [20]. To the best of our knowledge, this study represents the first metagenome-derived functional inference of the oyster microbiome.

The amplicon-based 16S rRNA gene profiling revealed low richness in oyster tissues as compared to the surrounding seawater (**Figure 1** and **Supplemental Figure S1**), while high variability was observed among different individuals and tissue types (**Supplemental Figure S2**). Statistically significant differences have been identified between the composition of gut and mantle communities as compared to other tissue types, while less support has been found for the distinctions among gill, hemolymph, pallial fluid, and inner shell swab samples (**Supplemental Figure S3**). The open circulatory system and absence of anatomical structures that would further compartmentalize organs, such as a serous membrane or peritoneal cavity, may explain the overlap observed in the community profiles among the gill, hemolymph, pallial fluid, and inner shell swab samples. These features, for example, have previously been speculated to play a role in the spread of *P. marinus* among oyster tissues [12]. Interestingly, the gill and mantle samples demonstrated significant differences in the beta diversity measurements despite their close contact in the oyster, indicating potential mechanistic differences in the association of microbes with oyster gill and mantle tissues.

Five major bacterial taxa have been identified at the class level, with significant differences in relative abundance across tissues observed for the *Gammaproteobacteria* (dominated by the *Vibrio* genus), *Spirochaetia*, *Mollicutes*, and *Chlamydiae*. Meanwhile, a more consistent relative abundance of *Fusobacteriia* was observed across the six different tissue types examined (**Figure 2**). The presence of these major taxa has been further confirmed with metagenomic sequencing of the oyster gut microbiome, where representative full-length 16S rRNA genes were reconstructed for each taxon, and six of the 16S rRNA gene sequences had identical matches to ASVs targeting the V4 region. Two of the metagenome-assembled 16S rRNA genes were classified as *Mollicutes* (*Mollicutes-1* and *Mollicutes-2*). While the *Mollicutes- 1* occurred primarily in the oyster gut and had a visibly higher but not statistically significant relative abundance in the uninfected than the *P. marinus* infected gut samples, the *Mollicutes-2* presented broadly among different tissue types and had a significantly higher relative abundance in the uninfected gill and pallial fluid samples compared to the *P. marinus* infected samples (**Supplemental Figure S5**). The differential abundance of Mollicutes sequences between uninfected and infected eastern oyster samples is similar to a prior study of the Sydney rock oyster (a Pacific species) where sequences related to *Mycoplasma* were present in the digestive gland of uninfected oysters but absent in oysters infected with the protozoan parasite, *Marteilia sydneyi* [24].

Additional insights into the oyster gut microbiome have been achieved with the reconstruction of MAGs. Overall, two MAGs of high completeness and low contamination have been identified for the *Mollicutes* and *Chlamydiae*, and one partial MAG has been identified from the *Spirochetes*. Specifically, phylogenomic assignment and the calculation of pairwise ANI and AAI values with reference genomes suggest the *Mollicutes* and *Chlamydiae* MAGs are distinct from existing isolates (**Supplemental Table S4, S5**). Genome-scale metabolic reconstruction of the *Mollicutes* MAG has revealed a heavy reliance on host-derived nutrients, with several unique metabolic pathways identified in the *Mollicutes* MAG as compared to other neighboring strains in the phylogeny. One is a chitin utilization pathway, which supports the degradation of chitin for carbon and energy metabolism; another is a complete ADI pathway that could fuel ATP production through the utilization of arginine, an abundant amino acid in the eastern oyster [43]. Arginine has been previously implicated in infection dynamics of oysters.

For example, *P. marinus* is speculated to sequester arginine to avoid host immune responses mediated by the production of nitric oxide, for which arginine is a precursor [44]. Interestingly, despite the presence of a GlcNAc utilization pathway in *M. marinum* and *M. todarodis* and the presence of an arginine deiminase gene in *M. mobile*, the complete ADI and chitin degradation pathways were only present in the *Mollicutes* MAG (**Figure 3B**).

The presence of the *Mollicutes* in the adult oyster gut has been observed in other 16S rRNA gene profiling studies [17,39,45]. A previous study also identified electron-dense bodies in eastern oyster gut goblet cells using transmission electron microscopy that resemble strains of the genus *Mycoplasma* [46]. Consistent with these prior studies, our analysis has provided additional support to the potentially universal presence of *Mollicutes* in the adult oyster gut. The absence of *Mollicutes* in adult biodeposits may also indicate a close association of the *Mollicutes* with digestive tissues, and this association may be mediated through a metabolic reliance of the *Mollicutes* on the host via the chitin utilization and arginine deiminase pathways (**Figure 3B**).

Interestingly, the *Mollicutes* were also largely absent from samples of homogenized eastern oyster larvae, which was in contrast to the higher observed relative abundance of the *Vibrio* genus (**Supplemental Figure S8**). In addition, sequencing of surrounding water samples of the adult oyster revealed a largely distant phylogenetic relationship between the water-associated and the oyster-associated *Mollicutes* populations (**Figure 4**), indicating a potential enrichment of specific *Mollicutes* strains in the host. Further study of the mechanisms of the *Mollicutes* acquisition, persistence, and physiology will begin to shed light on the nature of the relationship between the oyster host and *Mollicutes* (i.e. commensal, pathogen, mutualist). Overall, the integrated study of the taxonomic composition and metabolic potential of the eastern oyster microbiome will set the stage for future research on bacterial transmission dynamics, host range, and relative impacts on host health.

## MATERIALS AND METHODS

### Sample Collection and Dissection

Sixty live oysters were sampled from an aquaculture farm on Ninigret Pond, RI (USA) on 25 August 2017. Oysters were transported to the laboratory in coolers on ice. Upon arrival at the laboratory, the outer shells of individual oysters were scrubbed to remove visible sediments. For ten of the oysters, hemolymph and pallial fluid were collected by creating a small notch in the shell. Pallial fluid was drained through the notch prior to shucking, and the hemolymph was obtained with a sterile syringe from the pericardial cavity. Pallial fluid and hemolymph samples were centrifuged at 16,100 x g for 8 minutes, the supernatant removed, and the pellet was resuspended in 500 µL of RNAlater (Invitrogen, Carlsbad, CA). A sample of the gut, gills, mantle, and a swab of the inner shell (taken from the inner surface of the top and bottom shells using sterile cotton swabs) were stored individually in 1 mL of RNAlater (Invitrogen, Carlsbad, CA). A mixed sample of mantle, gut, and rectum tissue was also preserved for pathogen quantification. In addition, 1 L of surface water collected at Ninigret Pond was pre-filtered through a 153 µm mesh filter. Then, the water sample was filtered onto a 5 µm pore size (diameter of 47 mm) filter (Sterlitech, Kent, WA) followed by a 0.2 µm pore size Sterivex filter (Millipore, Burlington, MA). All samples were stored in a −80°C freezer before further processing. Detailed tracking of all samples and their application in the 16S rRNA community profiling and metagenomic sequencing is provided in **Supplemental Table S1**.

### Perkinsus marinus Quantification

For each of the 60 oysters, DNA from mantle, gut, and rectum tissue samples was extracted using a previously described protocol with modifications [47]. First, 5 mg of archived oyster tissue was sterilely placed into a 1.5 mL microcentrifuge tube with 160 µL of preheated urea buffer and incubated at 60°C in a dry bath for one hour. Samples were vortexed intermittently throughout the incubation. After incubation, the tubes were vortexed again and placed into a 95°C dry bath for 15 minutes. Samples underwent centrifugation for 5 minutes at 15,000 x g, and 100 µL of the supernatant was removed and mixed with 1 uL of 100X TE buffer, 50 µL ammonium acetate, and 400 µL of 95% ethanol.

After mixing, the samples were left at −20°C overnight to precipitate. The following day, samples underwent centrifugation for 20 minutes at 15,000 x g to pellet the precipitate and were washed three times with 70% ice-cold ethanol to remove any remaining impurities. DNA was eluted in 100 µL of buffer (10 mM Tris-HCl, 0.1 mM EDTA, pH 8.0). Quantity and quality of the DNA were assessed with the NanoDrop 2000c (Thermo Scientific), and the samples were diluted to 100 ng in 3 µL to ensure consistency between qPCR reactions.

Subsequently, the DNA was used as a template in a multiplex, real-time PCR reaction testing for *P. marinus*, MSX, and SSO. *P. marinus* primers and dual-labeled probes from [48] were used in conjunction with MSX/SSO primers, MSX probe, and SSO probe (primer and probe sequences in **Supplemental Table S8**). Each 20 µL reaction was carried out using 300 nM *P. marinus* primers, 450 nM MSX/SSO primers and 75 nM probe with 3.44 µL water, 10 µL iQ Multiplex Powermix (BioRad, Hercules, CA), and 1 µL template DNA. The thermal cycler protocol was as follows: 95°C for 60 seconds, followed by 40 cycles of 95°C for 15 seconds, then 60°C for 15 seconds.

### DNA Extraction of Individual Tissue Samples

DNA extractions were performed with the Qiagen Allprep PowerFecal DNA/RNA Kit (Qiagen, Hilden, Germany) following the manufacturer’s protocol with slight modifications. Modifications included the addition of 2 µL of Proteinase K (Fisher Scientific, Waltham, MA) and 13.5 µL of 20% SDS (Fisher Scientific, Waltham, MA) to the lysis buffer. Mechanical lysis was performed via two rounds of homogenization with a TissueLyser (Qiagen, Carlsbad, CA) for 2.5 minutes each at 30 Hz, and the samples were incubated for five minutes at room temperature in between.

### 16S rRNA Community Profiling

16S rRNA gene profiling was performed using template DNA amplified with V4 primers 515F/806R with Illumina adapter overhangs (**Supplemental Table S8**) and 2x Phusion HF Master Mix (Thermo Fisher Scientific, Waltham, MA) [49, 50]. The PCR was configured in a touchdown protocol and contained the following program: 3- minute initial denaturation at 94°C; followed by 10 cycles of a 45-second denaturation at 94°C, 60-second annealing at 60°C with every cycle decreasing 1°C, and 30 seconds of elongation at 72°C; followed by 25 cycles of a 45-second denaturation at 94°C, 60-second annealing at 50°C, and 30 seconds of elongation at 72°C; 10 minutes of final extension at 72°C, and an indefinite hold at 4°C. Sequencing libraries were prepared at the Rhode Island Genomics and Sequencing Center (RIGSC) using a standardized protocol as follows: 50 ng of Ampure XP cleaned PCR product was used as a template in a second PCR reaction (5 cycles) with 2x Phusion HF Master Mix in order to attach full indices and adapters with the Illumina Nextera Index Kit (Illumina, San Diego, CA). The resultant PCR products were cleaned with Ampure XP, profiled with an Agilent BioAnalyzer DNA1000 chip (Agilent Technologies, Santa Clara, CA), quantified using a Qubit fluorometer (Invitrogen, Carlsbad, CA), and normalized for pooling. Then, the pooled library was quantified with qPCR with a Roche LightCycler 480 using the KAPA Biosystems Illumina Kit (KAPA Biosystems, Woburn, MA). The final pooled library underwent 2×250 bp sequencing on an Illumina MiSeq (Illumina, San Diego, CA).

### Microbial Cell Enrichment of Pooled Gut Samples

In order to enhance the metagenomic binning of co-assembled contigs, a microbial cell enrichment procedure was performed on the gut samples through the adaptation of a procedure previously applied to marine sponges [51]. The gut samples of 10 oysters were pooled and homogenized with 10 ml of ice-cold, sterile artificial seawater (28 ppt) in an autoclaved mortar and pestle and vortexed for 10 minutes. The homogenized sample was centrifuged at 100 x g at 4°C for 30 minutes. The supernatant was extracted and centrifuged again at 10,967 x g at 4°C for 30 minutes. After removing the supernatant, the resultant pellet was washed three times with ice-cold, sterile artificial seawater. The pellet from the final round of centrifugation was diluted with ice-cold, sterile artificial seawater and pushed through a 3 µm pore size (diameter of 47 mm) filter (Millipore, Burlington, MA) with a syringe. The flowthrough was subsequently pushed through a 0.2 µm pore size (diameter of 22 mm) filter (Sterlitech, Kent, WA). DNA extraction was then performed on the 0.2 µm filter using the Qiagen QIAamp PowerFecal DNA Kit following manufacturer protocols. Mechanical lysis was performed via two rounds of homogenization with a TissueLyser (Qiagen, Carlsbad, CA) for 1 minute each at 30 Hz, and the sample was incubated for five minutes at room temperature in between.

### Shotgun Metagenomic Sequencing

Two rounds of metagenomic sequencing were performed. In the first round, 12 gut samples were selected from the samples in which 16S rRNA gene profiling was performed (**Supplemental Table S1**). In the second round, a single library was prepared using the microbial enriched DNA from pooled homogenate of oyster gut samples (described above). Preparation of metagenomic libraries was performed at the RIGSC using standardized protocols including DNA fragmentation with an S220 ultrasonicator (Covaris, Woburn, MA), quantification with a Qubit fluorometer (Invitrogen, Carlsbad, CA), library preparation (including end repair, A-tailing, adapter ligation, and size selection) on the Wafergen Apollo 324 with PrepX reagents and consumables (Takara Bio, Kusatsu, Japan), followed by quality control using an Agilent BioAnalyzer High Sensitivity chip (Agilent Technologies, Santa Clara, CA), and quantification of the final library with qPCR with a Roche LightCycler 480 using the KAPA Biosystems Illumina Kit (KAPA Biosystems, Woburn, MA). Sequencing of the metagenomic libraries was performed at Duke University’s Sequencing and Genomic Technologies Core. For the first round, the pooled library was sequenced 2×150 bp across three lanes on an Illumina HiSeq 4000, while the second round was sequenced 2×150 bp on an Illumina NextSeq 500 (Mid-output).

### Amplicon Sequence Analysis

Demultiplexed read pairs were analyzed using the Quantitative Insights Into Microbial Ecology 2 (QIIME2) software version 2018.4 [52]. Using the DADA2 plugin, samples were subjected to quality filtering, denoising, and chimera detection [53]. Following quality control and chimera removal of the sequencing data, samples with less than 5,000 total read pairs were dropped from further analysis due to the low sequencing depth (**Supplemental Table S2)**. Taxonomic assignment was performed with QIIME2’s classifier by mapping to the SILVA database release 132 [54]. Mitochondrial and chloroplast sequences were removed following the taxonomic assignment. The resulting feature table and taxonomy data were exported and analyzed in R version 3.6.0. Alpha and beta diversity analyses were performed on the feature table with the vegan package in R [55]. Relative abundance of a taxon was calculated through dividing the ASV counts of that taxon in a given sample by the total number of ASVs in the same sample. The raw reads from three previous eastern oyster microbiome studies were retrieved from the NCBI Sequence Read Archive (SRA) linked to the following BioProject accession numbers: PRJNA518081 [27], PRJNA504404 [40], and PRJNA386685 [39] and subjected to the same analyses as specified above. Based on metadata from the SRA, samples that were not from the eastern oyster (water, mussels, etc.) were removed from the analysis after denoising was completed.

### Metagenomic Assembly and Binning

The data from both rounds of metagenomic sequencing underwent trimming and adapter removal with Trimmomatic (leading and trailing bases below a quality score of 3 were removed, a 4 bp sliding window was used with an average quality score of 15, and a minimum read length of 130 bp was required) and the last 10bp of all reads were removed with the CutAdapt software [56, 57]. Due to differences in the chemistry associated with the Illumina NextSeq and HiSeq platforms, the reads from the two rounds of metagenomic sequencing underwent slightly different quality control measures. The reads from the sample that was sequenced on the Illumina NextSeq 500 also underwent additional steps in Trimmomatic and CutAdapt in order to remove PolyG. In Trimmomatic, a sequence composed of 50 guanine residues was added to the adapter sequences, and a sequence of 10 guanine residues was used as an adapter sequence in CutAdapt (allowing for an error rate of 20%).

The quality filtered reads were then mapped with BBMap to a combined reference database containing the *C. virginica* genome (RefSeq Assembly Accession GCF_002022765.2), the *P. marinus* genome (RefSeq Assembly Accession GCF_000006405.1), all complete bacterial genomes downloaded on 18 April 2018, and a collection of manually curated draft genomes of bacteria isolated from oysters (strain names and NCBI RefSeq accessions in **Supplemental Table S9**). Reads mapped to a bacterial genome or unmapped to any genomes in the reference database were used for the metagenomic co-assembly and binning. Paired reads were considered as being mapped to a bacterial genome if at least one read from the pair was mapped. Metagenomic co-assembly was performed using MEGAHIT with default parameters [58]. Binning of the co-assembled contigs was performed with default parameters in MetaBat2 [59] using read mappings performed with BBMap, with the resulting sam files converted to bam format and sorted with SAMtools [60]. Completeness and contamination of the MAGs reconstructed from the oyster microbiome were assessed using CheckM. The *ssu_finder* function in CheckM was also used to identify 16S rRNA genes in the metagenomic co-assembly [61]. Candidate 16S rRNA genes greater than 1,000 bp in length were retained.

### Phylogenetic Positioning of *Mollicutes* ASVs

The ASVs taxonomically assigned to the class *Mollicutes* from this and prior studies [39], two full-length *Mollicutes* 16S rRNA genes assembled from the metagenomes collected in this study, and the full-length 16S rRNA genes obtained from four *Firmicutes* outgroups (**Supplemental Table S7**), were placed into a reference tree with SILVA’s ACT server, using the SINA aligner [62]. The minimum identity was set at 85% and the number of neighbors per query set at 100. The resultant phylogeny was collapsed based on the environment where the reference SILVA sequences were derived in iTol [63].

### Phylogenomic Analyses of *Mollicutes* and *Chlamydiae* MAGs

Phylogenomic reconstructions were performed on the *Mollicutes* MAG and the *Chlamydiae* MAG with reference genomes from the two corresponding taxa (**Supplemental Tables S4**, **S5**). To ensure consistency in gene calling, all genomes were first analyzed using Prodigal with genetic code 4 for the Mollicutes [64] and standard genetic code for the *Chlamydiae*. Conserved single-copy genes (CSCGs) were identified through the analysis of bidirectional best BLAST hits as described in a previous study [65]. The identified CSCGs were individually aligned with MUSCLE, the alignments were concatenated, and a phylogeny was reconstructed with RAxML using the JTT substitution model and the GAMMA model of rate heterogeneity. Pairwise average nucleotide identity (ANI) values were computed with OrthANI [66]. The average amino acid identity (AAI) was computed by the mean protein identity values of all bidirectional best BLAST hits identified based on the following thresholds: e-value less than or equal to 1×10^-3^; sequence identity greater than or equal to 30%; coverage of 70% or higher to both sequences in the alignment.

### Metabolic Reconstruction of the *Mollicutes* MAG

A metabolic reconstruction of the *Mollicutes* MAG was developed based on an initial mapping of the proteome to the KEGG database through the KAAS server [67] and the annotation of transporters was based on homology searches to the Transporter Classification Database [68]. The draft reconstruction underwent manual curations through comparisons to three *Mycoplasma* isolates closely related to the *Mollicutes* MAG (i.e. *M. mobile*, *M. todarodis*, and *M. marinum*) and using reference annotations from an existing metabolic model of *Mycoplasma pneumoniae* [69]. The metabolic reconstruction was incorporated into a genome-scale model using PSAMM [70]. Metabolic pathway gaps were identified using the PSAMM *gapcheck* function, which in turn were used to guide the manual curation of gene functions from the MAG and from other contigs identified from the metagenomic co-assembly. For example, the pyruvate dehydrogenase complex was initially not identified in the *Mollicutes* MAG, but its subunits were found on other contigs in the co-assembly with top BLAST hits (in the NCBI non-redundant protein database) to members of the *Mycoplasma* genus. In this case, the pyruvate dehydrogenase reaction was included in the metabolic model to highlight the potential presence of this function. Some gap reactions, such as the transport of acetate, were included in the metabolic network to represent the potential export of acetate as a metabolic product.

## DATA AVAILABILITY

Raw sequencing data is available from the NCBI Sequence Read Archive under the BioProject accession PRJNA658576.

## ACKNOWLEDGEMENTS

We would like to thank Alexa Sterling, Ashley Hamilton, Benjamin Korry, Dina Proestou, Evelyn Takyi, Kathryn Markey Lundgren, Rebecca Stevick, and Samuel Hughes for advice and/or assistance related to sampling and processing of oysters. Thanks to Janet Atoyan of the Rhode Island Genomics and Sequencing Center for advice and assistance related to sample preparation for sequencing.

## AUTHOR CONTRIBUTIONS

Research Design & Planning: YZ, ZTP, MGC; Sample Collection: ZTP, MGC, KDT, YZ; *P. marinus* Quantification: AKS, RMS; Molecular Sequencing: ZTP, YZ; Data Analysis: ZP, YZ; Metabolic Reconstruction: KDT, KTR, ZTP, YZ; Manuscript Writing: ZP, YZ; Manuscript Editing and Approval: All authors.

## SUPPLEMENTAL FIGURES

**Figure S1.**
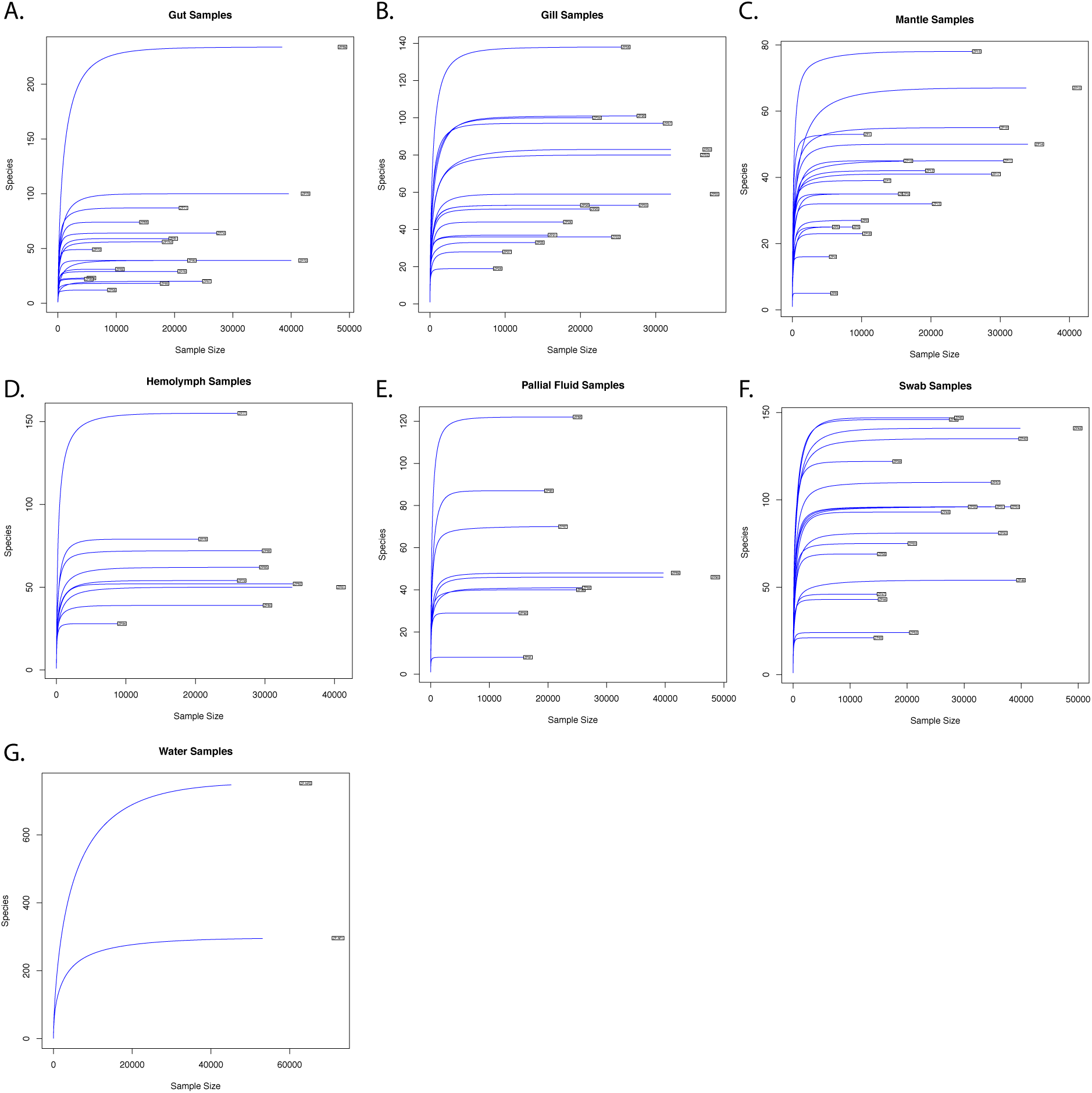
Rarefaction curves for the gut (**A**), gill (**B**), mantle (**C**), hemolymph (**D**), pallial fluid (**E**), inner shell swab (**F**), and water samples (**G**).

**Figure S2.**
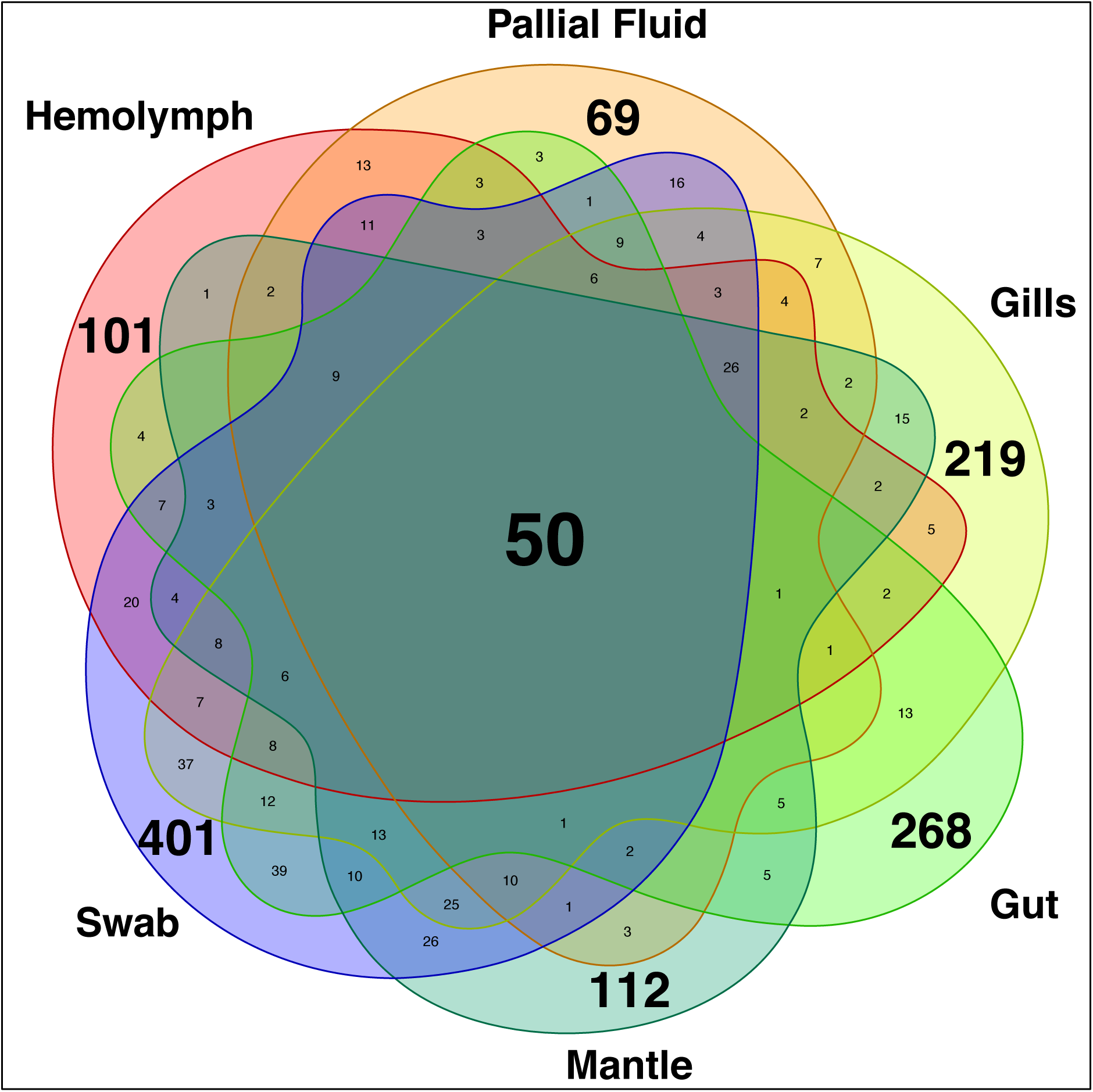
Venn diagram showing the overlap of ASVs identified from different oyster tissues. ASVs observed in at least one sample of a tissue type were counted as present in that tissue type. The number of samples in each tissue type can be found in **Supplemental Table S2**.

**Figure S3.**
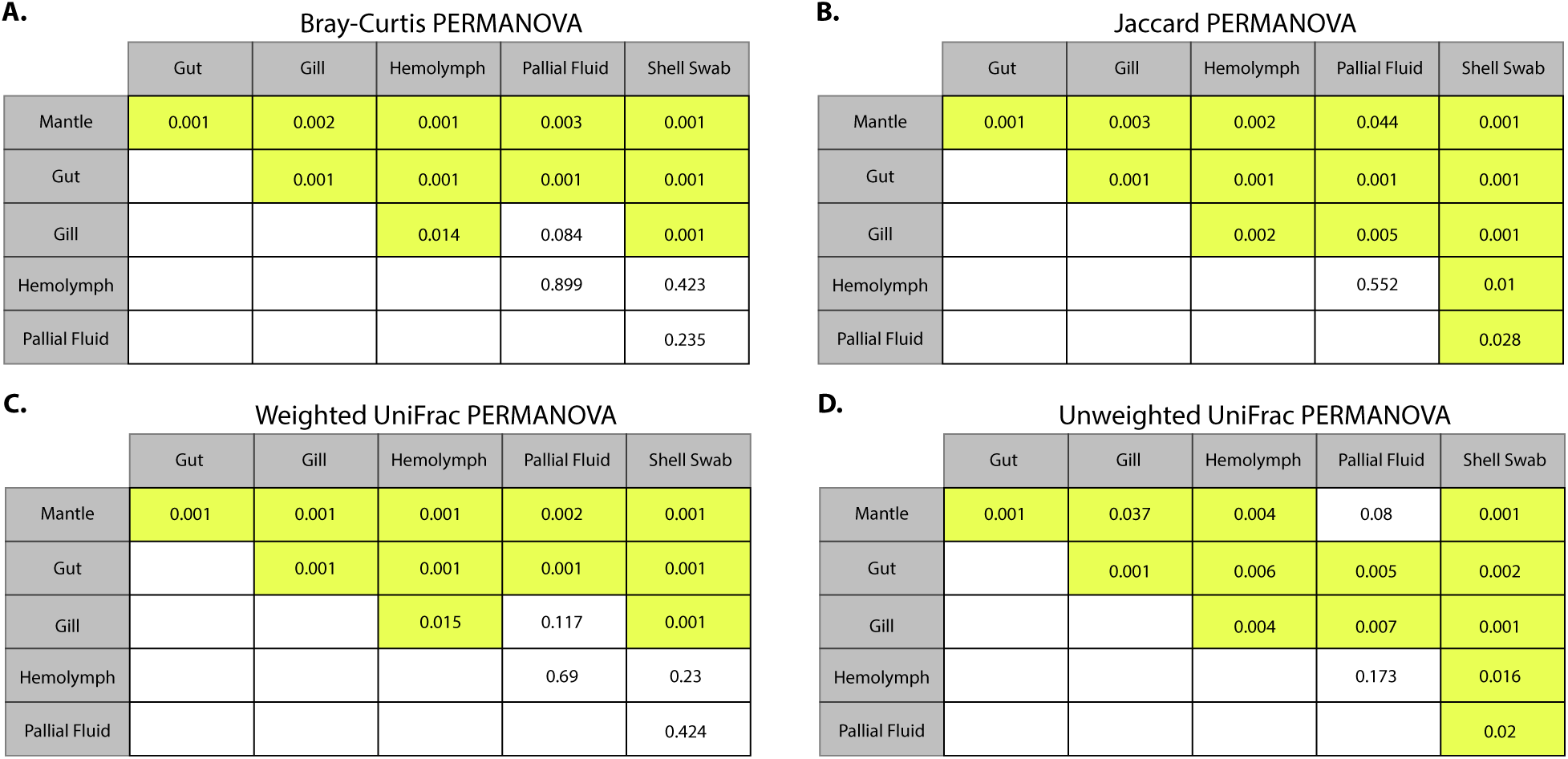
Statistical significance in the comparison of beta diversity between tissues based on four distance metrics: Bray-Curtis (**A**), Jaccard (**B**), Weighted UniFrac (**C**), and Unweighted UniFrac distances (**D**). Statistically significant differences were marked with a yellow background to the corresponding p-values based on the permutational multivariate analysis of variance (PERMANOVA) test (p-value < 0.05).

**Figure S4.**
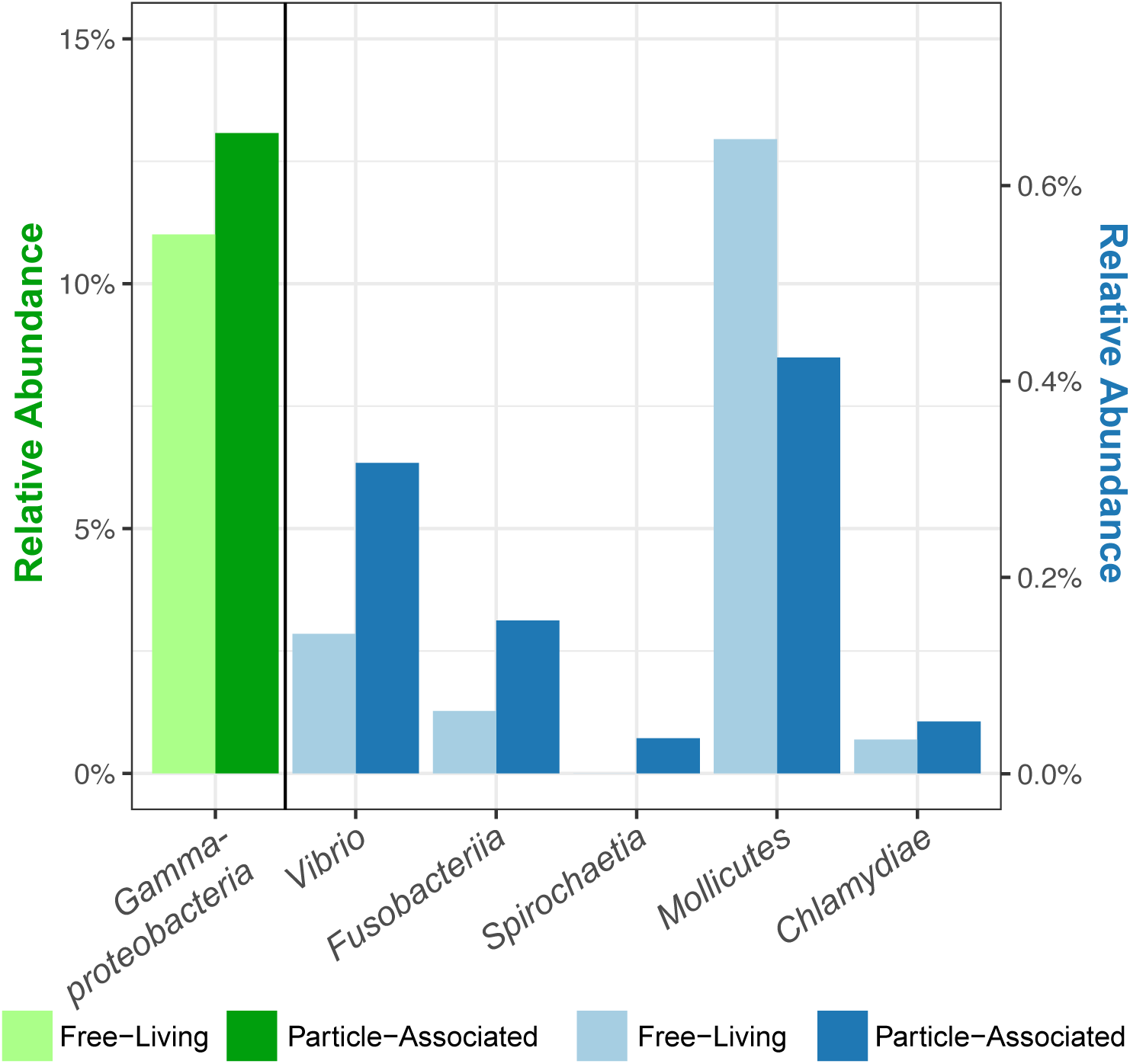
Relative abundance of the major taxa (*Gammaproteobacteria*/*Vibrio*, *Fusobacteriia*, *Spirochaetia*, *Mollicutes*, and *Chlamydiae*) in the free-living fraction (0.2-5 µm) and the particle-associated fraction (5-153 µm) of seawater samples. The relative abundance of the *Gammaproteobacteria* (in green), given their high value, are assigned to the primary (left) y-axis. The other taxa (in blue), are assigned to the secondary (right) y-axis.

**Figure S5.**
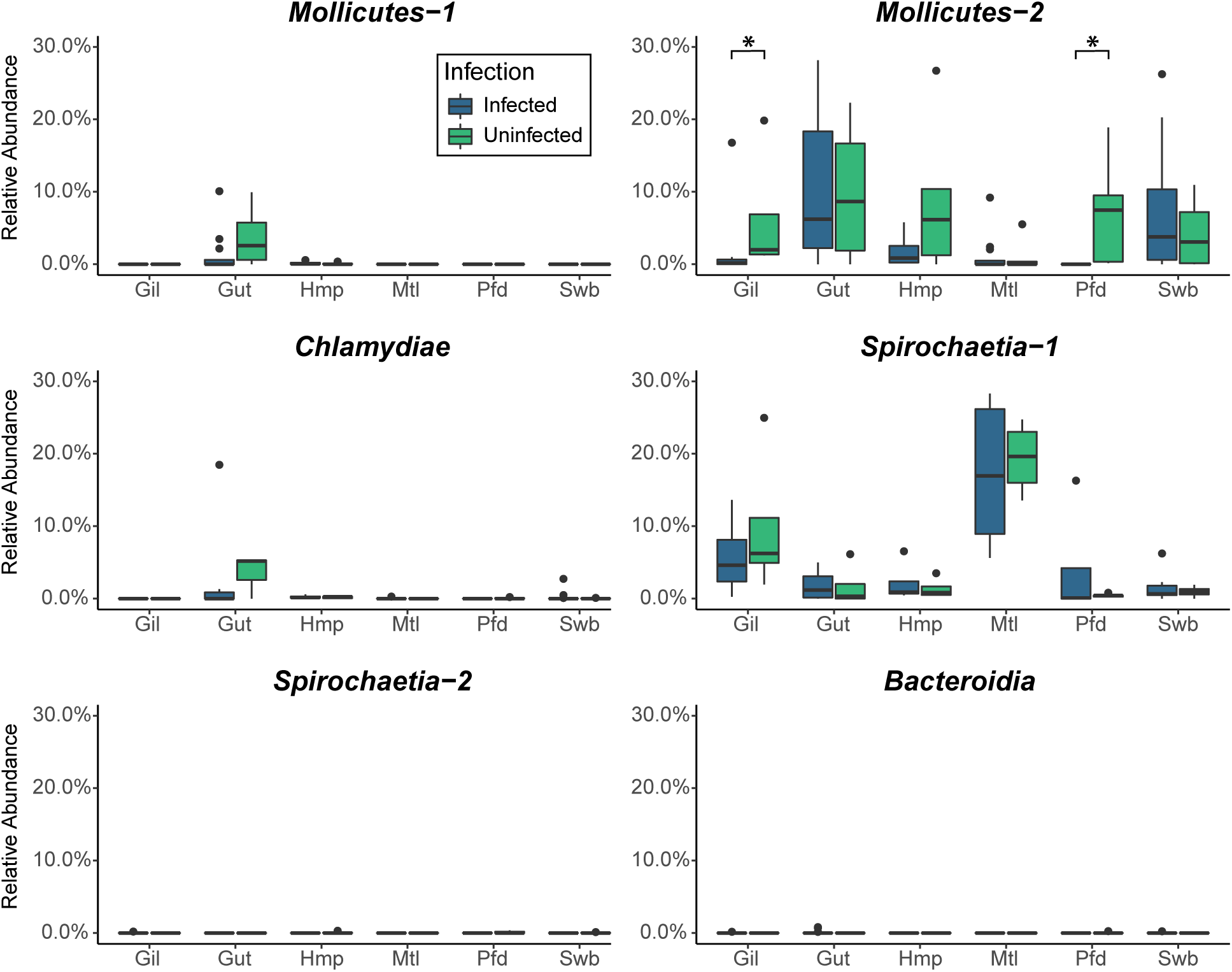
Relative abundance of 16S rRNA gene sequences from the metagenomic assembly as measured by their corresponding ASVs (**Supplemental Table S3**). The abundance measurements from different sample types were grouped based on the detection of *Perkinsus marinus* infections from individual oysters. Tissues that showed a significant difference between uninfected and *P. marinus* infected samples were marked: * p-value < 0.05.

**Figure S6.**
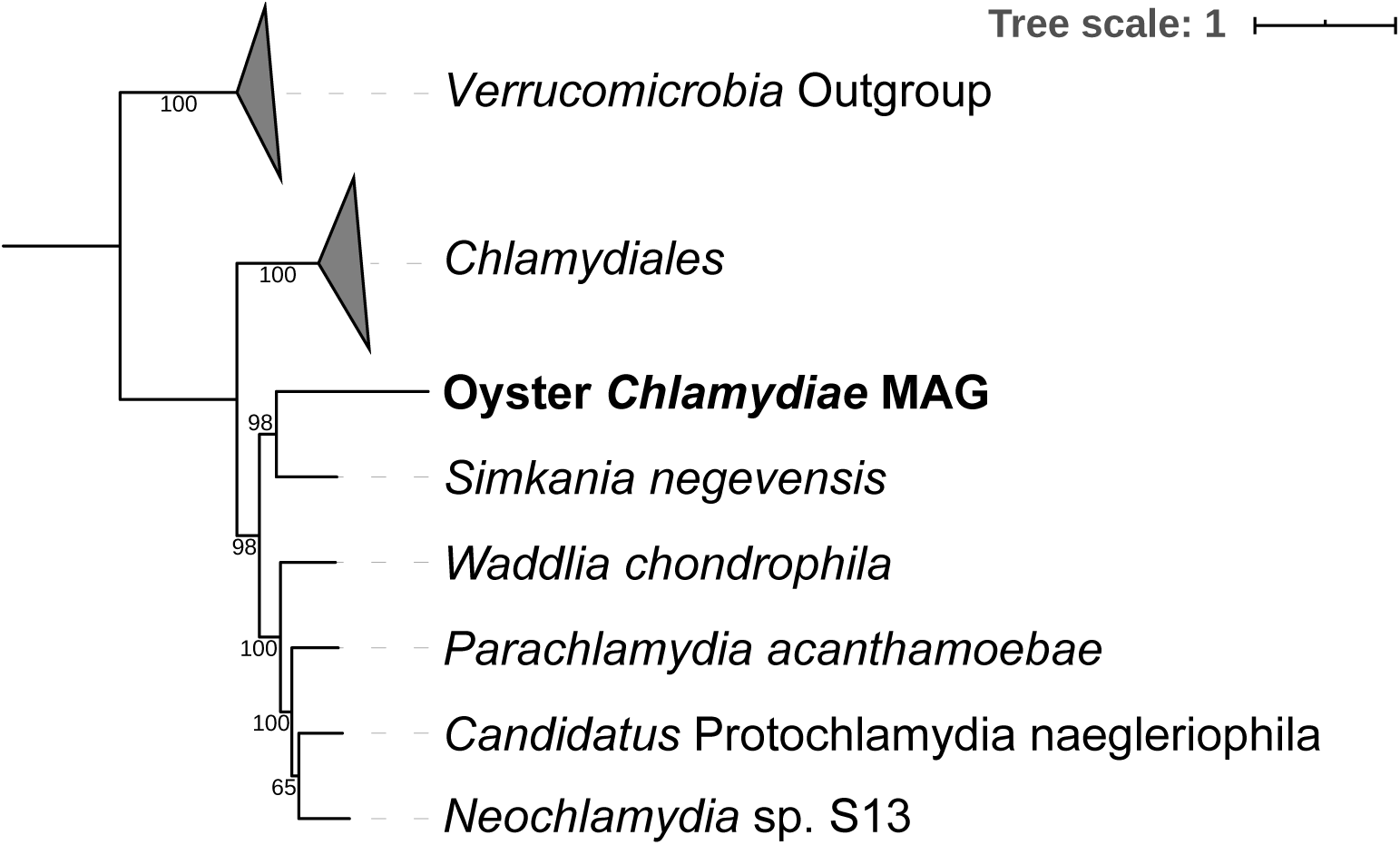
Maximum likelihood phylogenomic reconstruction of the oyster *Chlamydiae* MAG based on conserved single copy genes identified from the MAG and reference *Chlamydiae* genomes (**Supplemental Table S5**). Species of the *Chlamydiales* order were collapsed into a single clade and formed a neighboring clade to the *Parachlamydiales* clade that includes the oyster *Chlamydiae* MAG. Members of the *Verrucomicrobia* were used as outgroups in the phylogeny. Support values were based on 100 iterations of bootstrapping.

**Figure S7.**
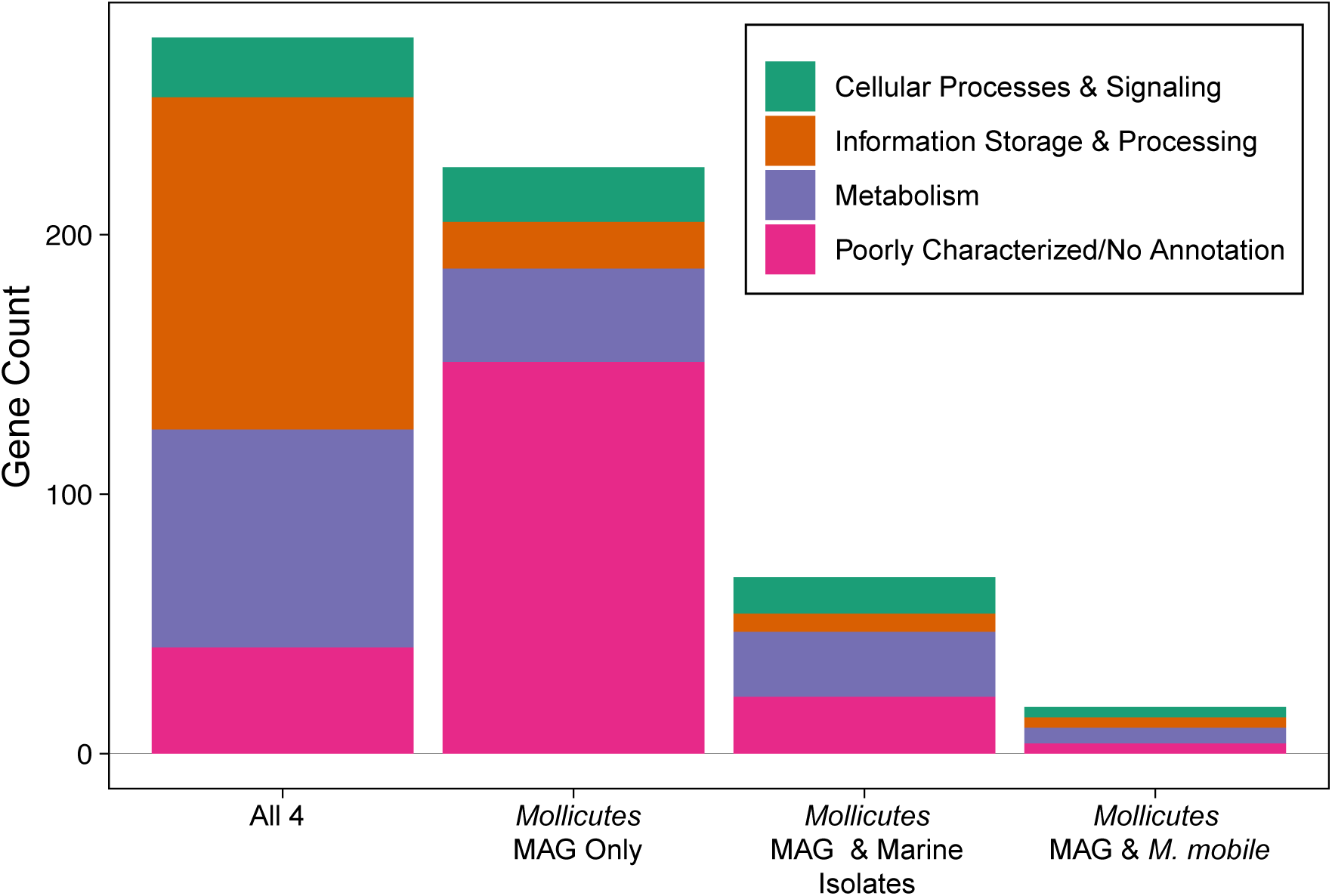
Distribution of COG functions among the conserved and unique genes of the oyster *Mollicutes* MAG. The COG functions were profiled among four distinct gene groups based on conservation between the oyster *Mollicutes* MAG and reference genomes, including the marine host-associated *Mycoplasma marinum* and *Mycoplasma todarodis*, and the freshwater host-associated *Mycoplasma mobile*: All 4–conserved in all four genomes; *Mollicutes* MAG Only– unique functions in the *Mollicutes* MAG; *Mollicutes* MAG & Marine Isolates–conserved between the *Mollicutes* MAG and the marine *Mycoplasma* (*M. marinum* and *M. todarodis*) but absent in *M. mobile*; *Mollicutes* MAG & *M. mobile*–conserved between the *Mollicutes* MAG and the freshwater *M. mobile*, but absent from the marine *Mycoplasma*.

**Figure S8.**
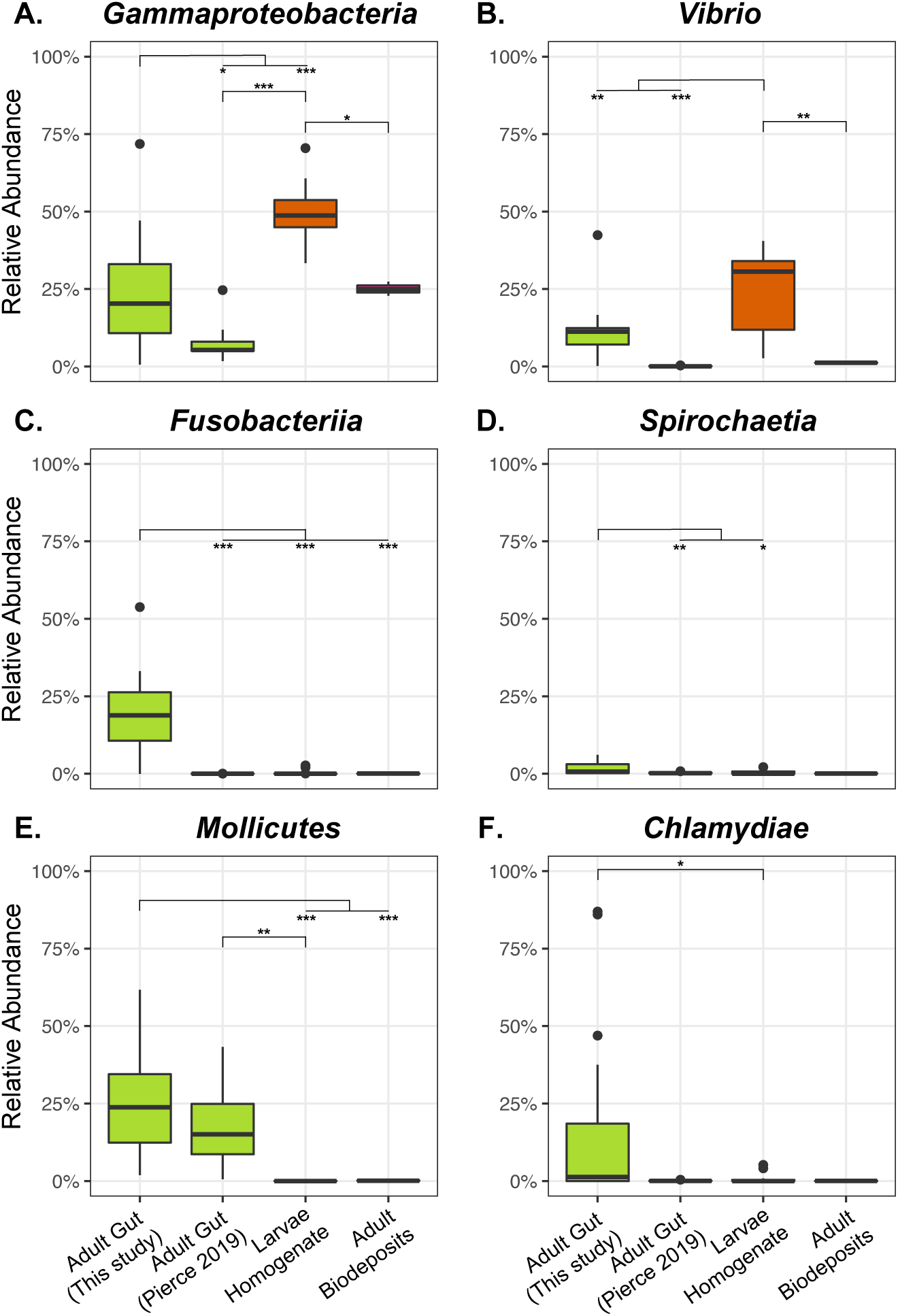
Comparison of relative abundances of the major taxa among different eastern oyster microbiome studies. Four distinct eastern oyster microbiome datasets were included in the comparison, including the adult gut samples from this study and from [39], larvae homogenate samples [27], and biodeposit samples [40]. Pairwise statistical significance was assessed with Tukey’s Honest Significant Differences test. Significant tissue pairs are marked as follows: * p- value < 0.05, ** p-value < 0.01, *** p-value < 0.001.

## SUPPLEMENTAL TABLES

**Table S1.**
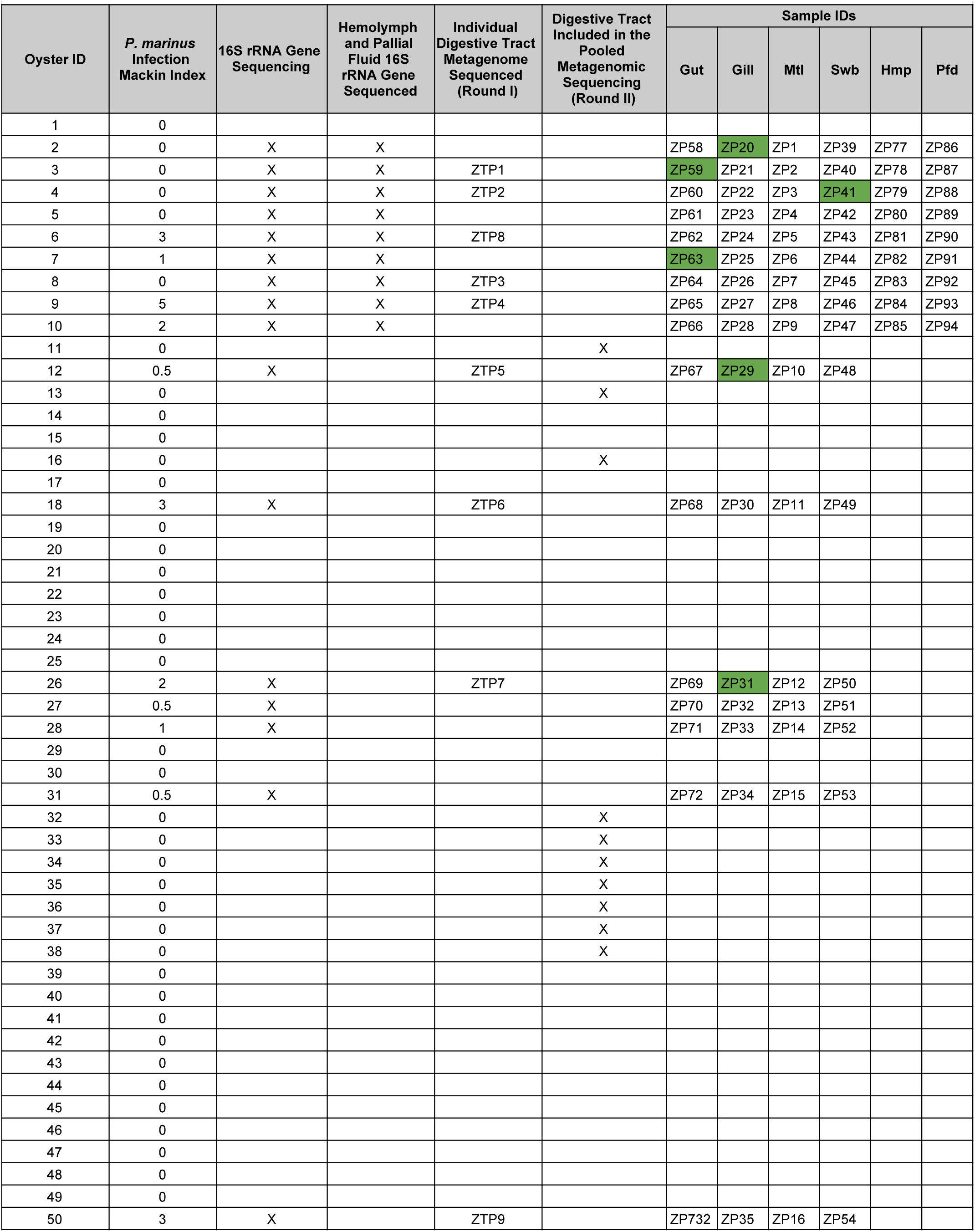
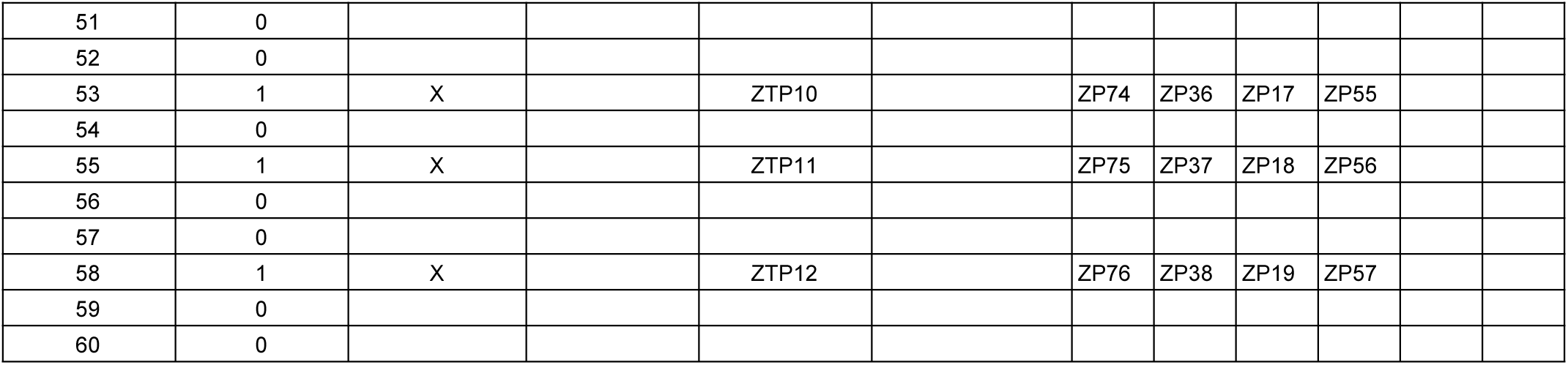
Summary of the oyster samples analyzed in this study. Sample identifiers for the amplicon sequencing were shown for each tissue sample of each individual oyster. Samples marked with a green background were removed from further analysis due to low sequencing depth.

**Table S2.** Number of samples obtained from different oyster tissues (before and after removing samples based on quality control) and summary statistics of the ASVs identified in each tissue type.

| Sample Type | Samples Sequenced | Samples Retained After QC | Median ASV Count Per Sample | 25th Percentile ASV Count Per Sample | 75th Percentile ASV Count Per Sample | Percentage of ASVs in > 80% Samples | % ASVs That Overlap With Water Samples |
| --- | --- | --- | --- | --- | --- | --- | --- |
| Gut | 19 | 17 | 39 | 23 | 64 | 1.65% | 9.90% |
| Gill | 19 | 16 | 53 | 36.75 | 86.5 | 2.78% | 10.90% |
| Mantle | 19 | 19 | 39 | 26 | 47.5 | 1.45% | 12.50% |
| Shell Swab | 19 | 18 | 94.5 | 57.75 | 119 | 1.29% | 10.00% |
| Pallial Fluid | 9 | 9 | 46 | 40 | 70 | 3.44% | 22.50% |
| Hemolymph | 9 | 9 | 54 | 50 | 72 | 5.77% | 18.90% |

**Table S3.**
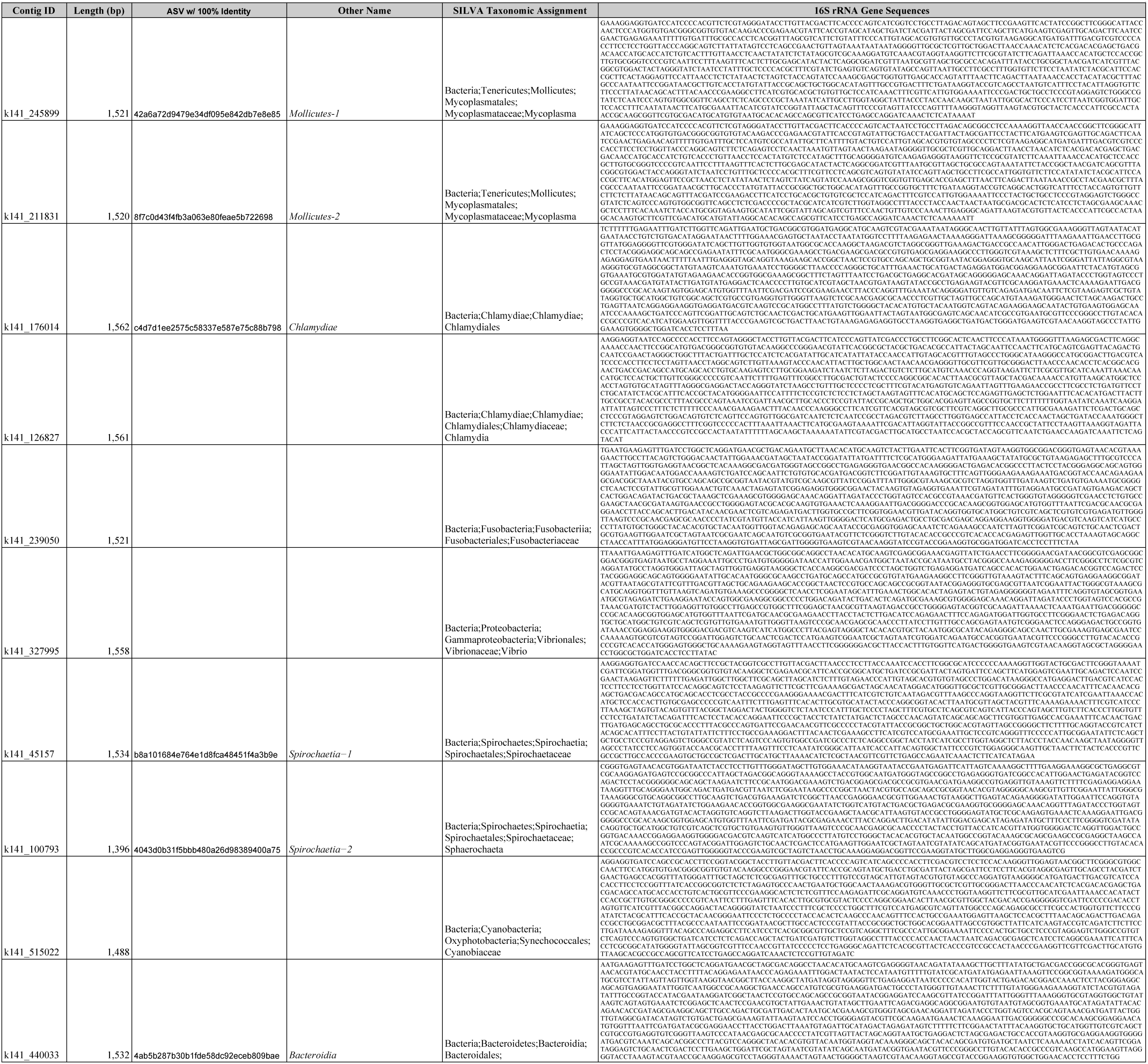
Summary of the metagenome-assembled full-length 16S rRNA genes. Six of the gene sequences were mapped to ASV sequences with 100% identity throughout the entire length of the ASVs. Taxonomic assignments were shown based on comparison to reference genes in the SILVA database.

**Table S4.**
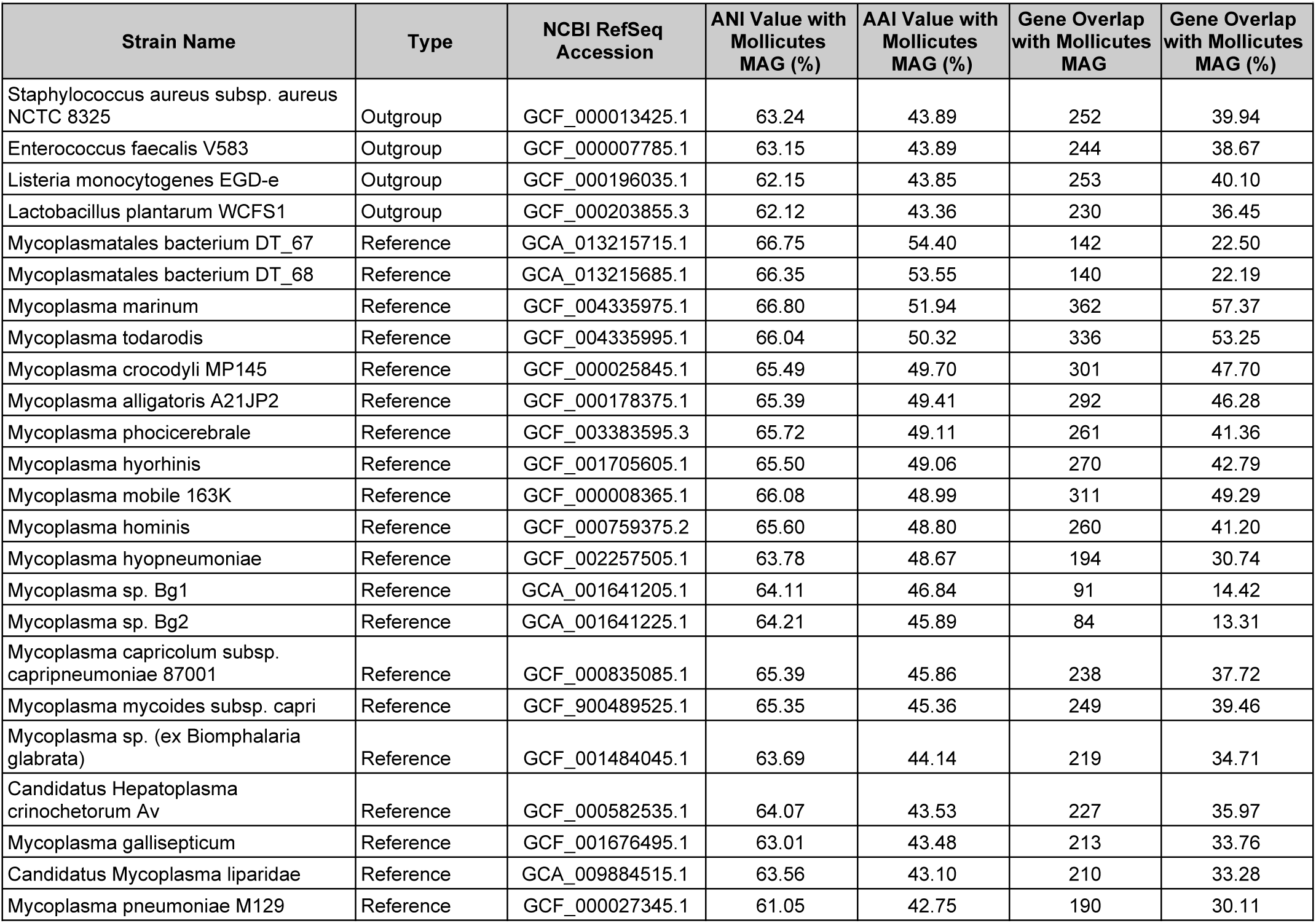
Average nucleotide identity (ANI) and average amino acid identity (AAI) of the oyster *Mollicutes* MAG compared to reference genomes and outgroups analyzed in the phylogenomic reconstruction (**Figure 3A**).

| Strain Name | Type | NCBI RefSeq Accession | ANI Value with *Mollicutes* MAG (%) | AAI Value with *Mollicutes* MAG (%) | Gene Overlap with *Mollicutes* MAG | Gene Overlap with *Mollicutes* MAG (%) |
| --- | --- | --- | --- | --- | --- | --- |
| *Staphylococcus aureus* subsp. *aureus* NCTC 8325 | Outgroup | GCF_000013425.1 | 63.24 | 43.89 | 252 | 39.94 |
| *Enterococcus faecalis* V583 | Outgroup | GCF_000007785.1 | 63.15 | 43.89 | 244 | 38.67 |
| *Listeria monocytogenes* EGD-e | Outgroup | GCF_000196035.1 | 62.15 | 43.85 | 253 | 40.10 |
| *Lactobacillus plantarum* WCFS1 | Outgroup | GCF_000203855.3 | 62.12 | 43.36 | 230 | 36.45 |
| *Mycoplasmatales bacterium* DT_67 | Reference | GCA_013215715.1 | 66.75 | 54.40 | 142 | 22.50 |
| *Mycoplasmatales bacterium* DT_68 | Reference | GCA_013215685.1 | 66.35 | 53.55 | 140 | 22.19 |
| *Mycoplasma marinum* | Reference | GCF_004335975.1 | 66.80 | 51.94 | 362 | 57.37 |
| *Mycoplasma todarodis* | Reference | GCF_004335995.1 | 66.04 | 50.32 | 336 | 53.25 |
| *Mycoplasma crocodyli* MP145 | Reference | GCF_000025845.1 | 65.49 | 49.70 | 301 | 47.70 |
| *Mycoplasma alligatoris* A21JP2 | Reference | GCF_000178375.1 | 65.39 | 49.41 | 292 | 46.28 |
| *Mycoplasma phocicerebrale* | Reference | GCF_003383595.3 | 65.72 | 49.11 | 261 | 41.36 |
| *Mycoplasma hyorhinis* | Reference | GCF_001705605.1 | 65.50 | 49.06 | 270 | 42.79 |
| *Mycoplasma mobile* 163K | Reference | GCF_000008365.1 | 66.08 | 48.99 | 311 | 49.29 |
| *Mycoplasma hominis* | Reference | GCF_000759375.2 | 65.60 | 48.80 | 260 | 41.20 |
| *Mycoplasma hyopneumoniae* | Reference | GCF_002257505.1 | 63.78 | 48.67 | 194 | 30.74 |
| *Mycoplasma* sp. Bg1 | Reference | GCA_001641205.1 | 64.11 | 46.84 | 91 | 14.42 |
| *Mycoplasma* sp. Bg2 | Reference | GCA_001641225.1 | 64.21 | 45.89 | 84 | 13.31 |
| *Mycoplasma capricolum* subsp. *capripneumoniae* 87001 | Reference | GCF_000835085.1 | 65.39 | 45.86 | 238 | 37.72 |
| *Mycoplasma mycoides* subsp. *capri* | Reference | GCF_900489525.1 | 65.35 | 45.36 | 249 | 39.46 |
| *Mycoplasma* sp. (ex *Biomphalaria glabrata*) | Reference | GCF_001484045.1 | 63.69 | 44.14 | 219 | 34.71 |
| *Candidatus Hepatoplasma crinochetorum* Av | Reference | GCF_000582535.1 | 64.07 | 43.53 | 227 | 35.97 |
| *Mycoplasma gallisepticum* | Reference | GCF_001676495.1 | 63.01 | 43.48 | 213 | 33.76 |
| *Candidatus Mycoplasma liparidae* | Reference | GCA_009884515.1 | 63.56 | 43.10 | 210 | 33.28 |
| *Mycoplasma pneumoniae* M129 | Reference | GCF_000027345.1 | 61.05 | 42.75 | 190 | 30.11 |

**Table S5.**
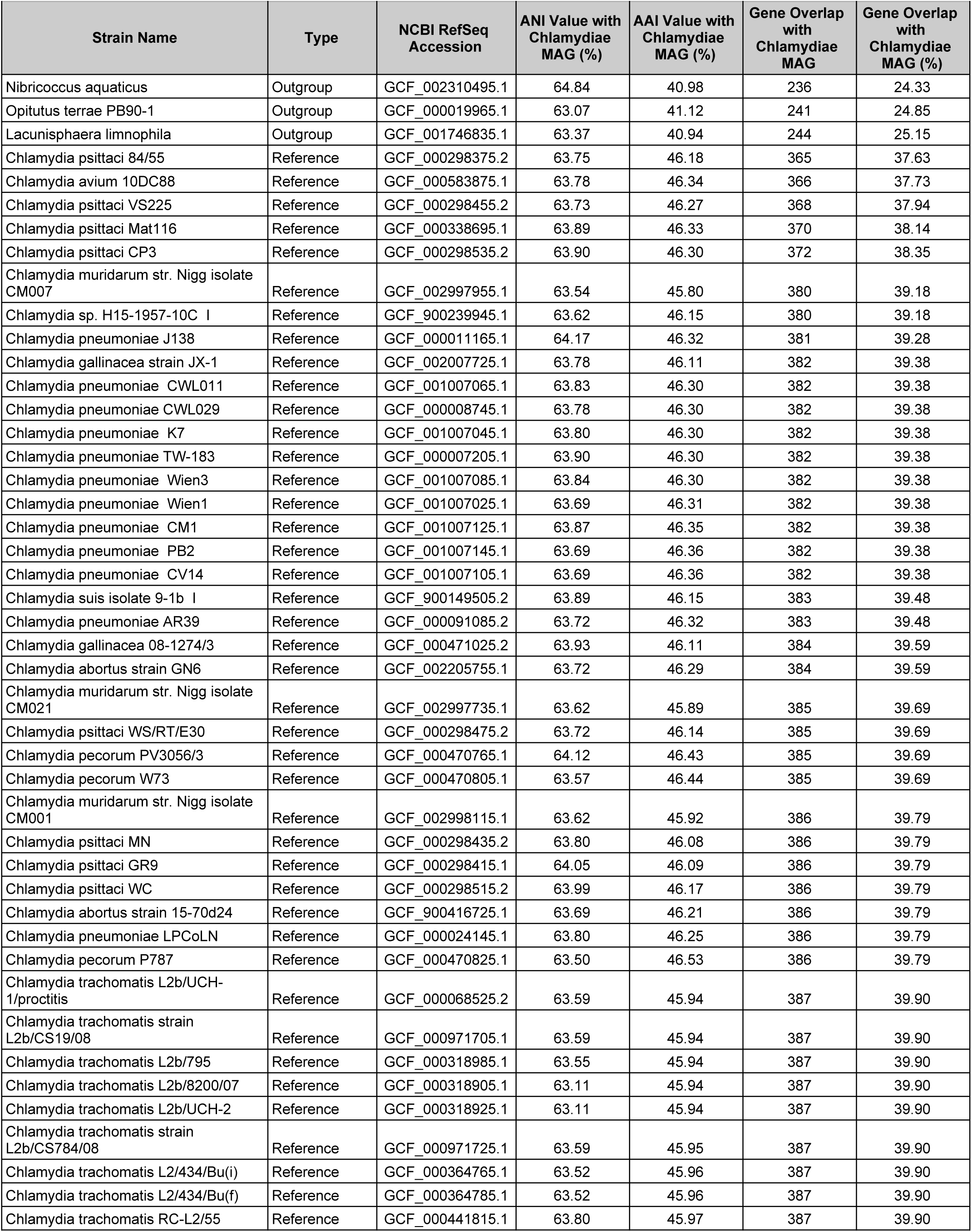
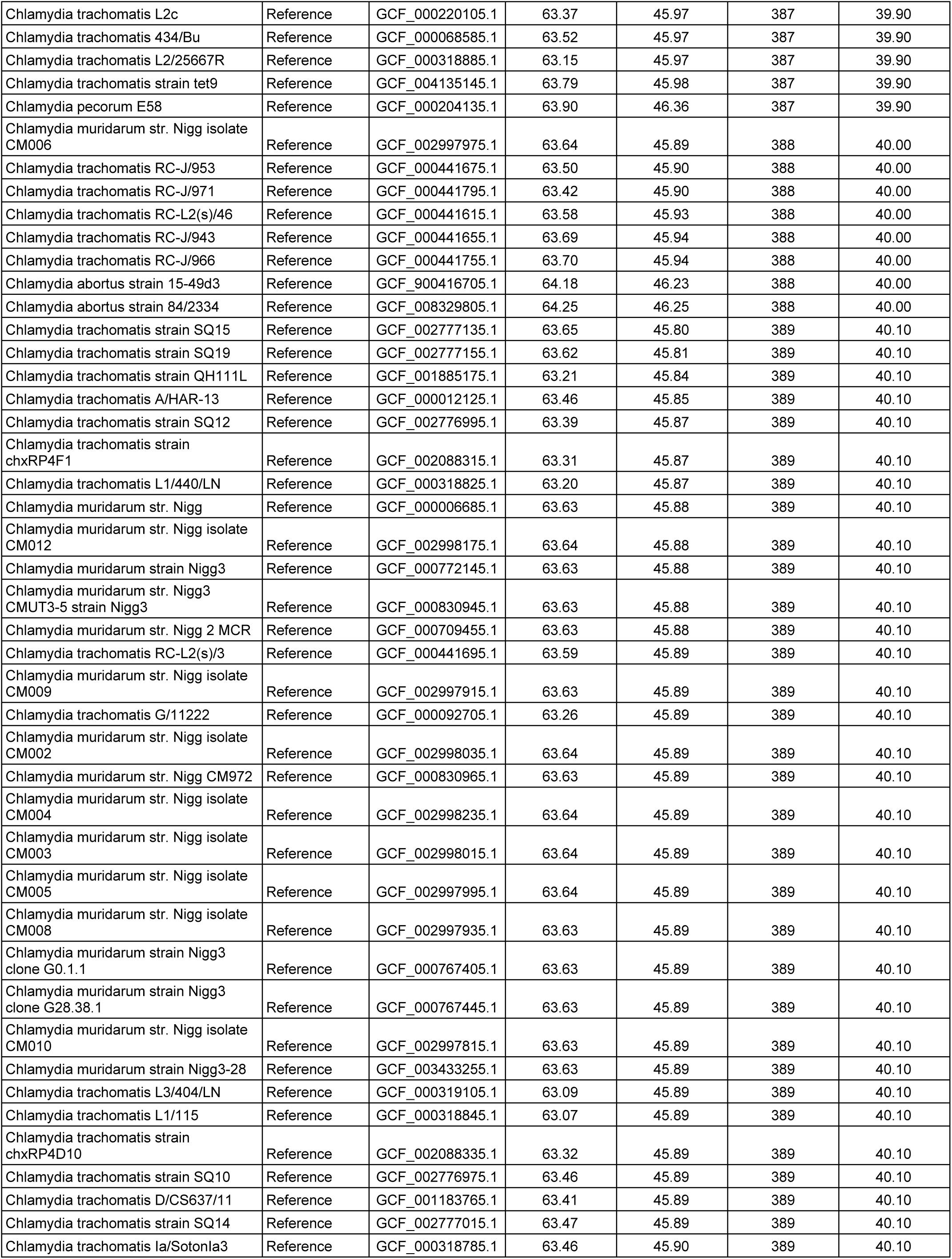
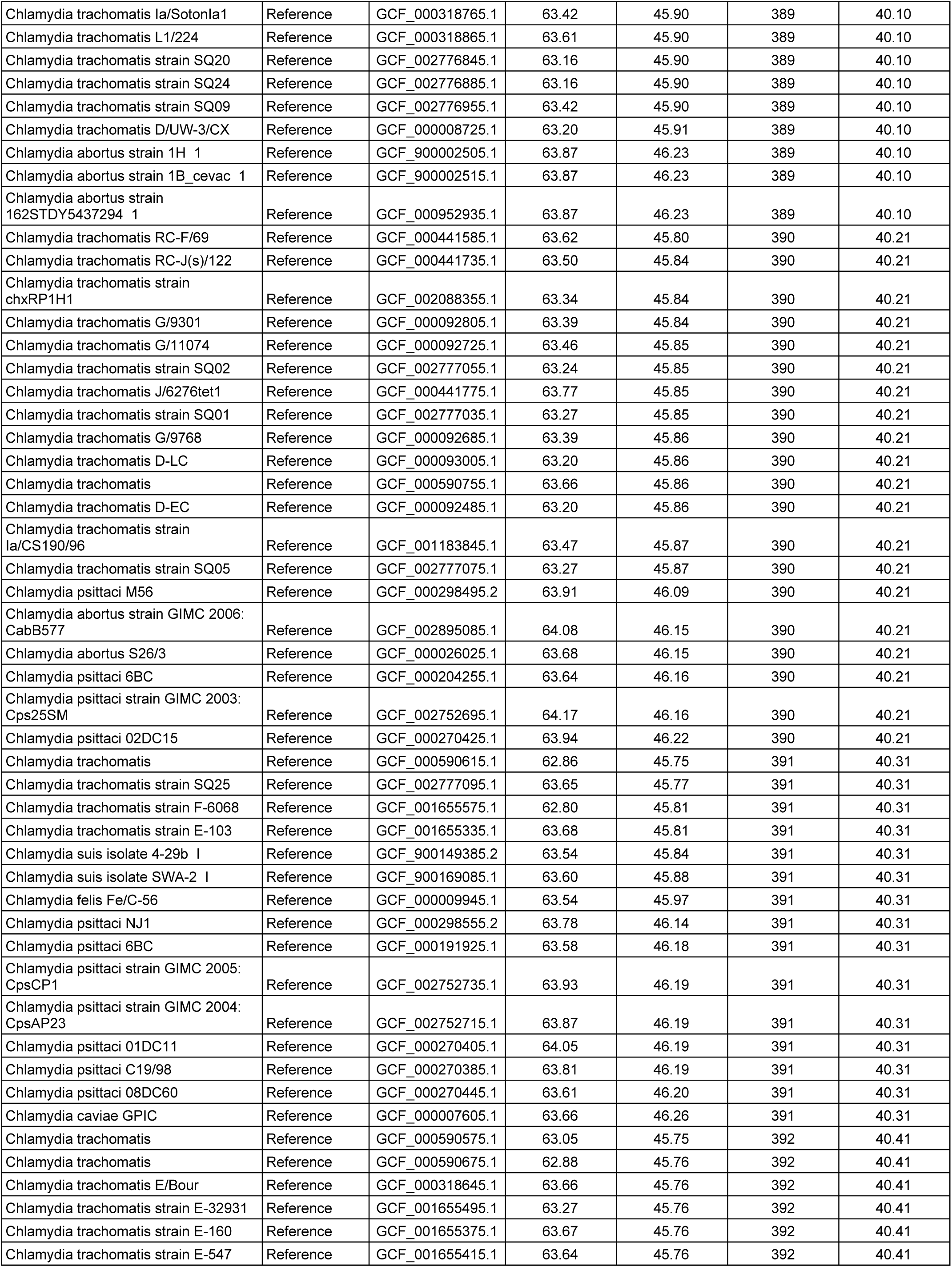
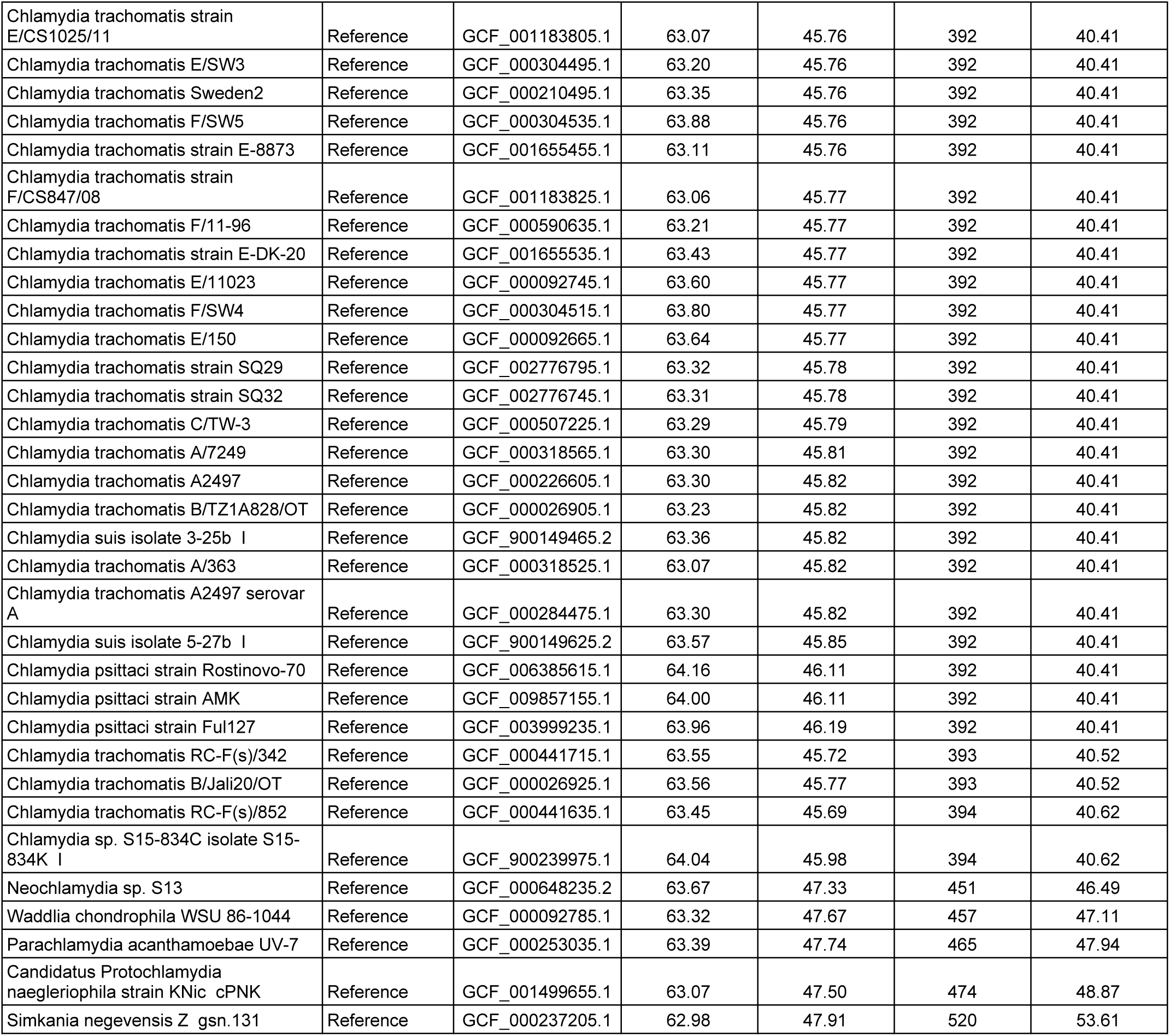
Average nucleotide identity (ANI) and average amino acid identity (AAI) of the oyster *Chlamydiae* MAG compared to reference genomes and outgroups analyzed in the phylogenomic reconstruction (**Supplemental Figure S6**).

**Table S6.**
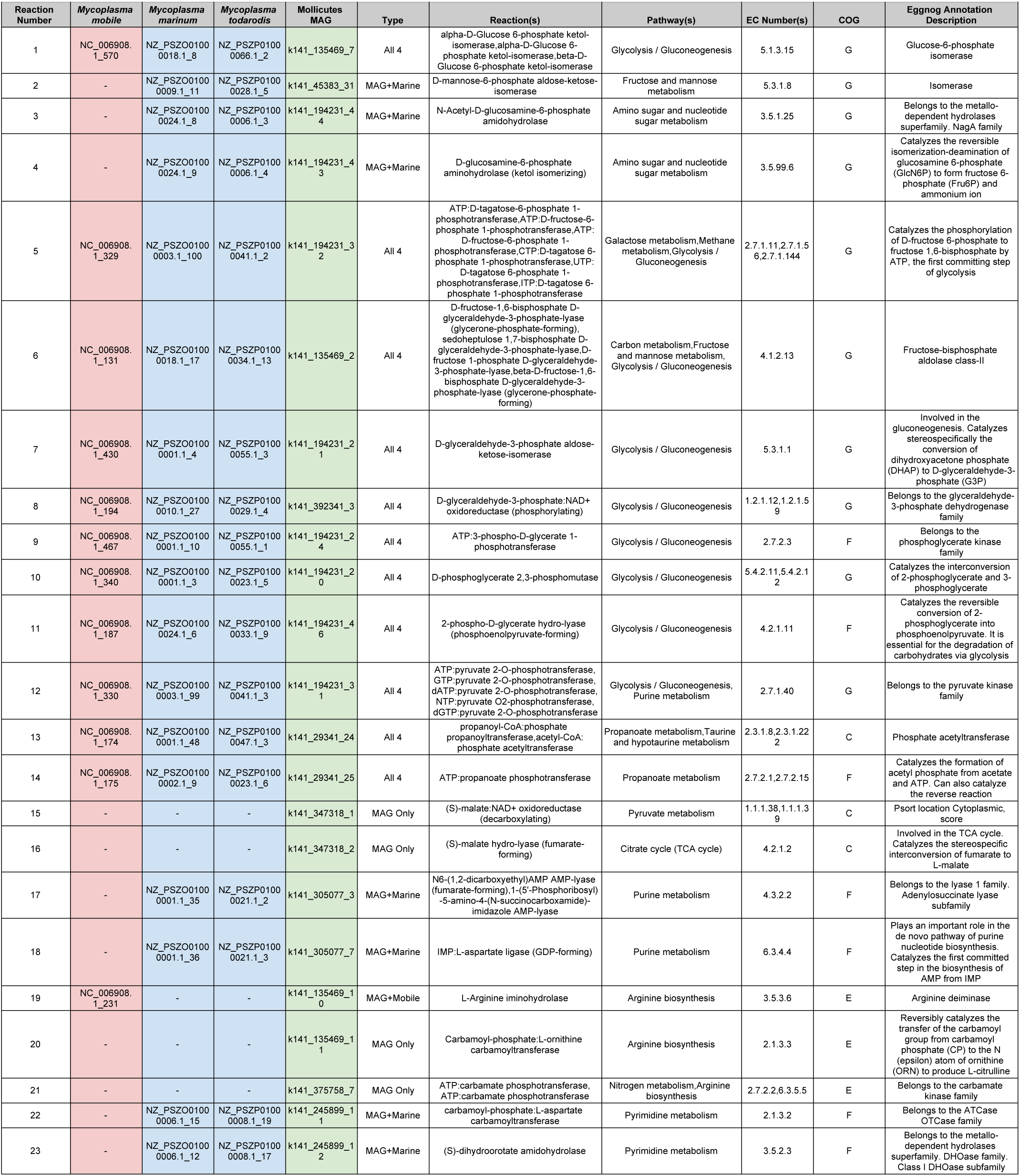
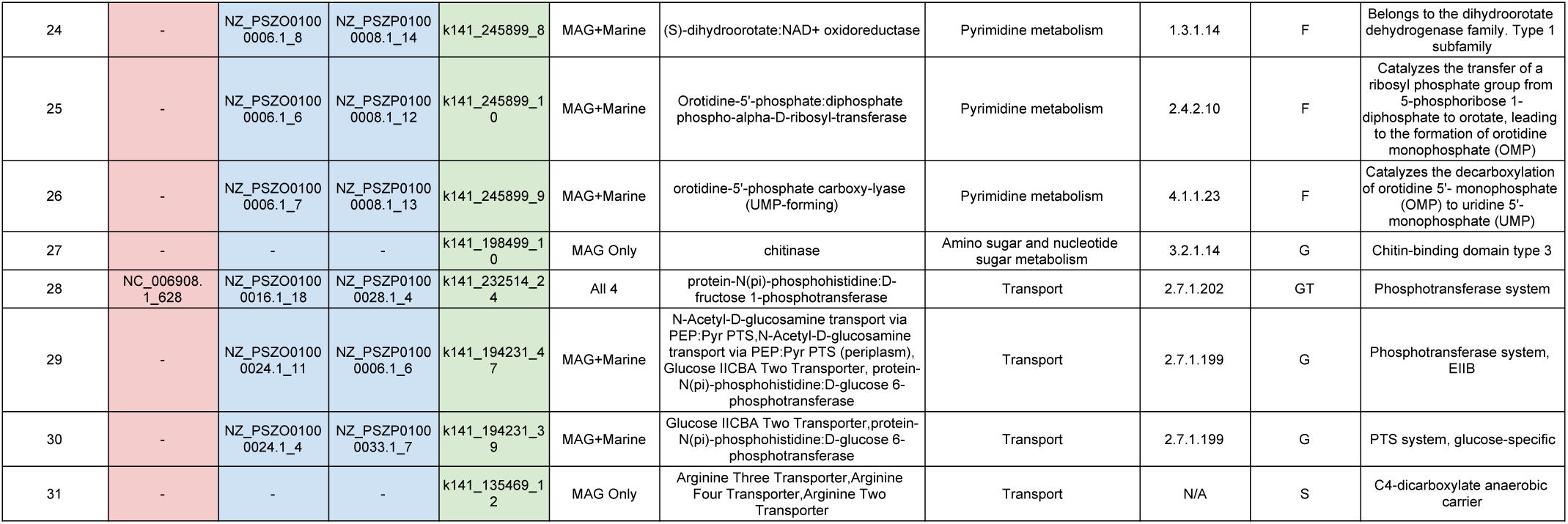
Metabolic functions corresponding to the visualization of central metabolism reconstructed from the oyster *Mollicutes* MAG. Each number in the Reaction Number column refers to a metabolic function correspondingly labeled in **Figure 3**. Gene identifiers corresponding to each function were included under the four genomes: *M. mobile*, *M. marinum*, *M. todarodis*, and oyster *Mollicutes* MAG. Each entry was classified in the Type column based on conservation of genes between the *Mollicutes* MAG and other genomes: All4–functions conserved in all four genomes analyzed; MAG+Marine–functions conserved between the MAG and the marine species *M. marinum* and *M. todarodis*; MAG+Mobile–functions conserved between MAG and *M. mobile*; MAG Only–functions unique to the MAG. Additional information includes the reaction name, metabolic pathway(s), Enzyme Commission Numbers (EC Number), COG classification, and eggNOG annotations were provided.

**Table S7.**
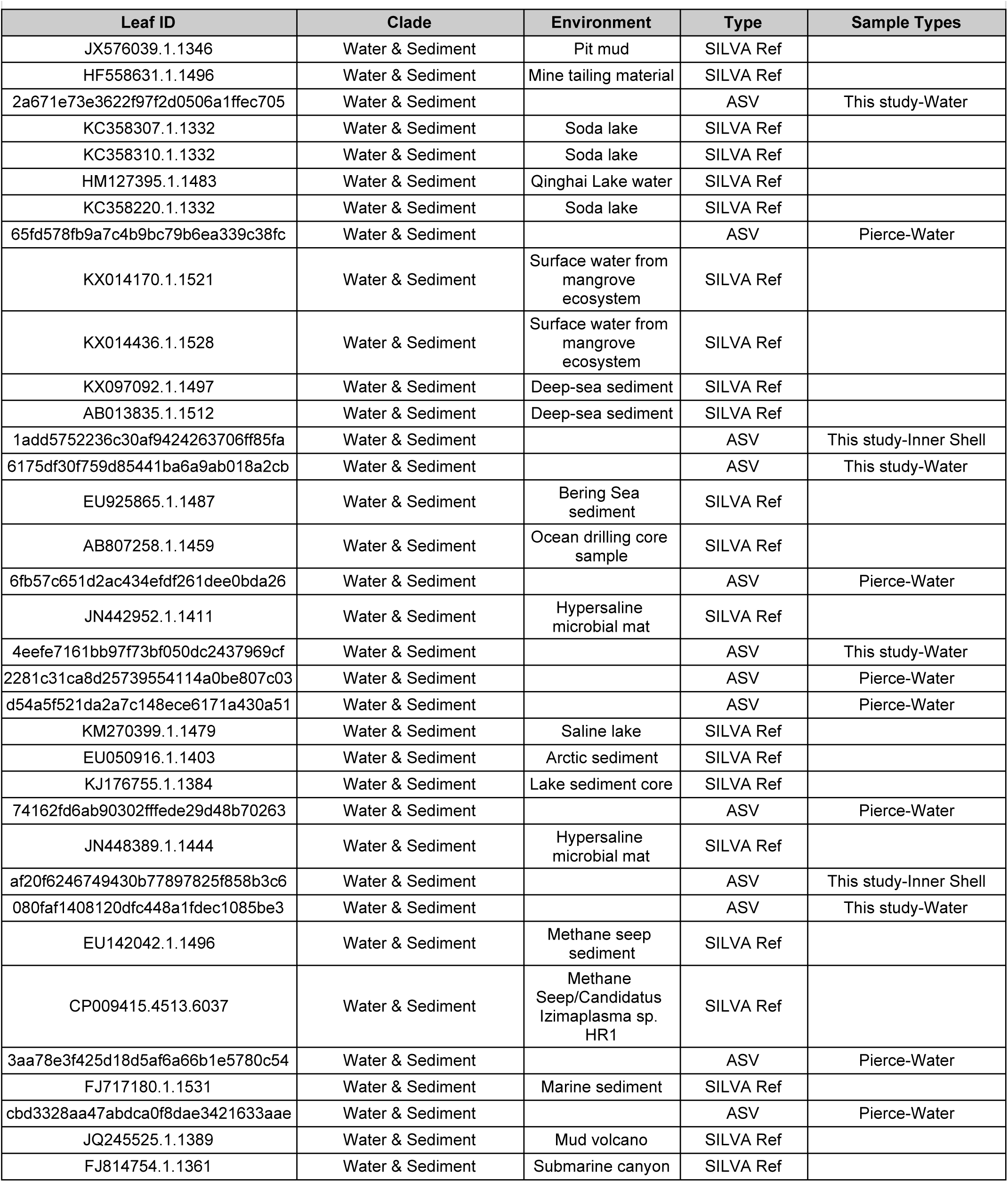
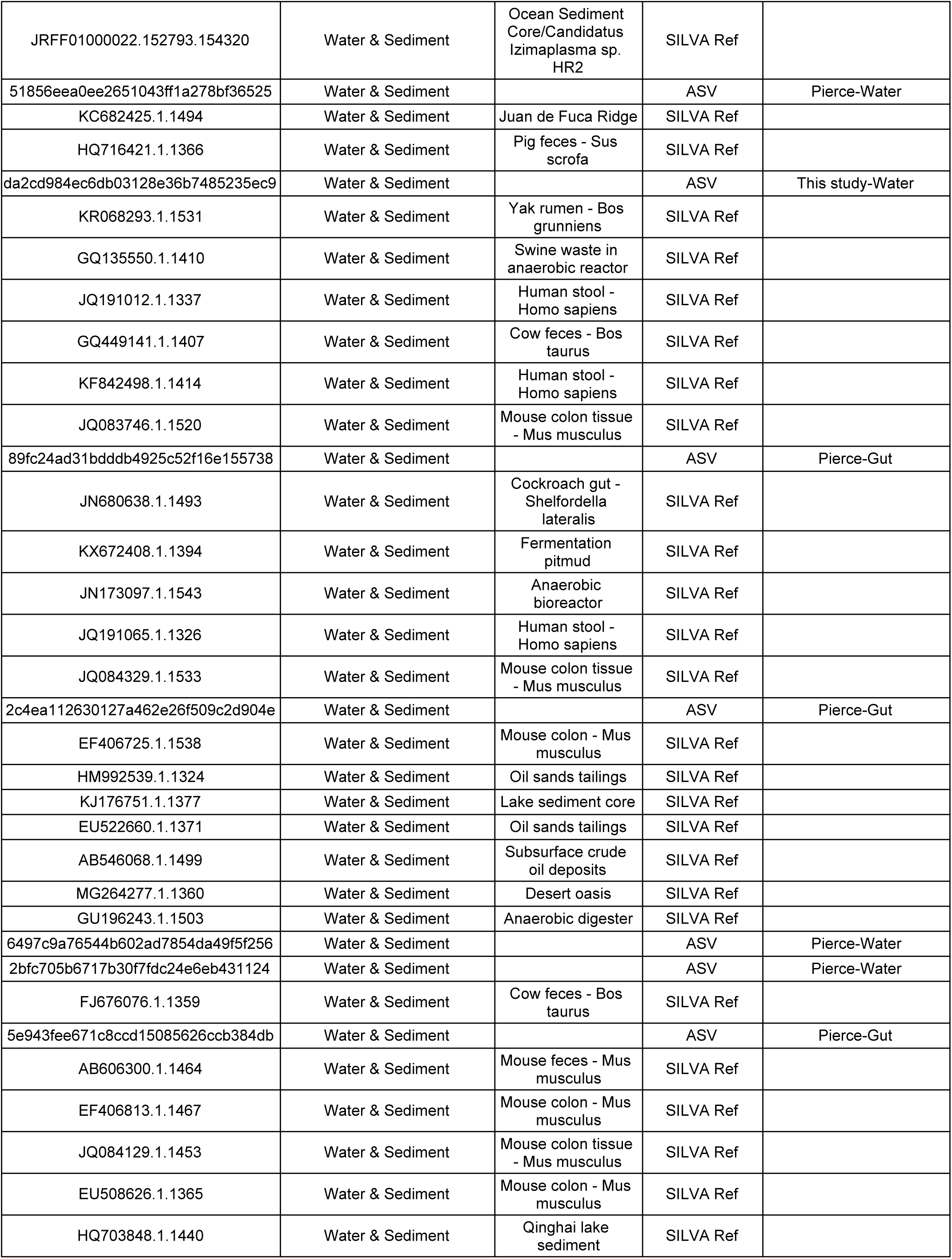
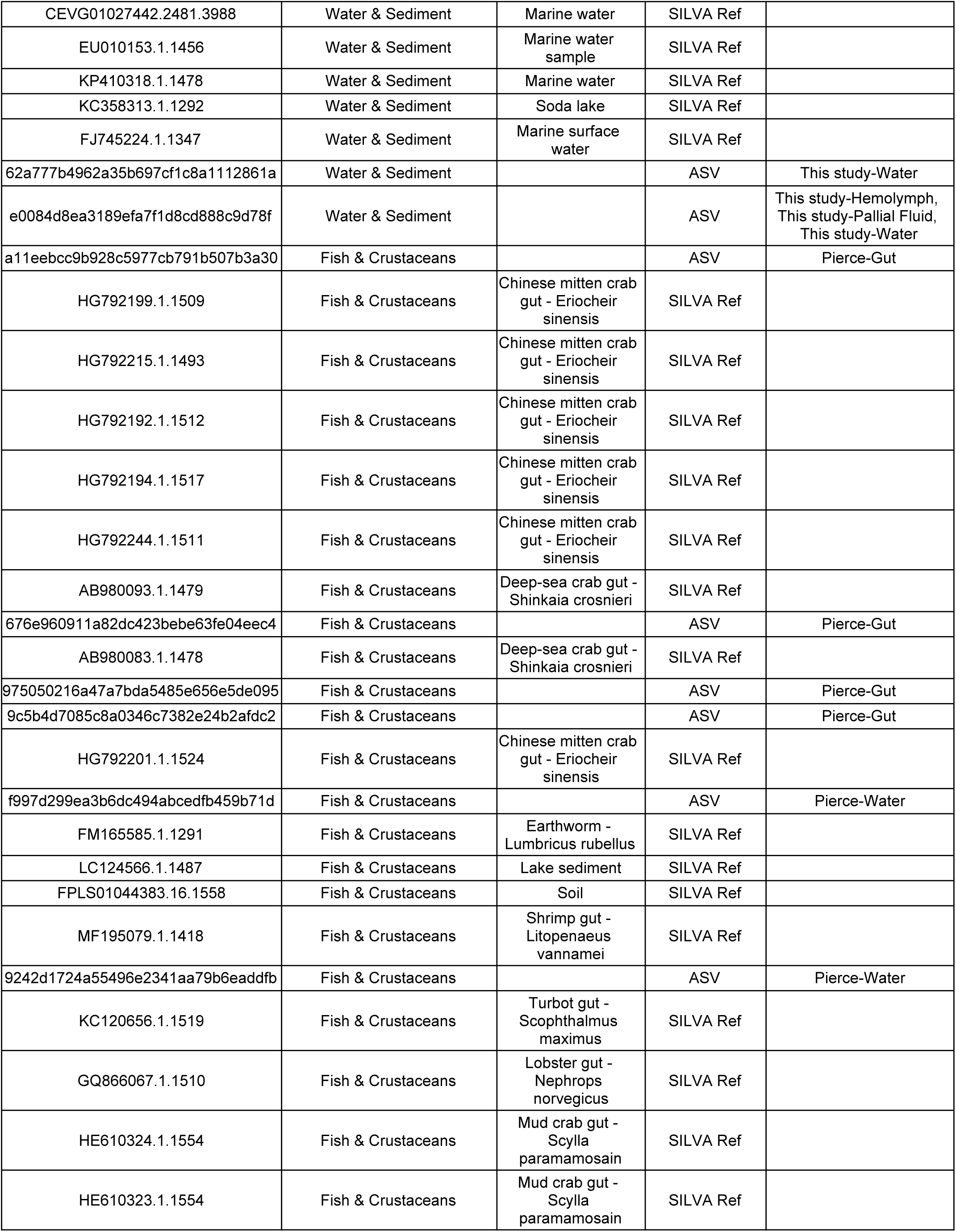
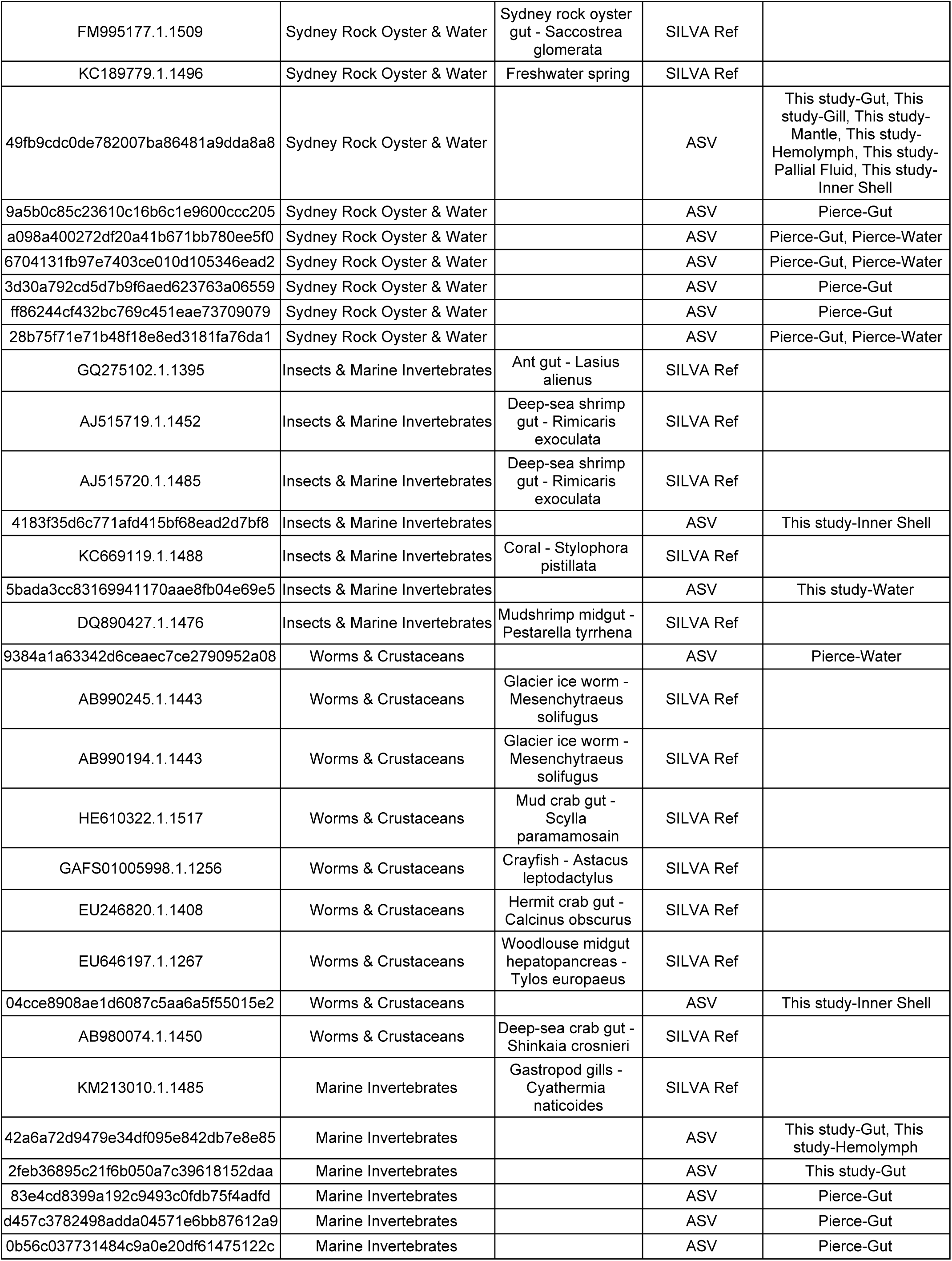
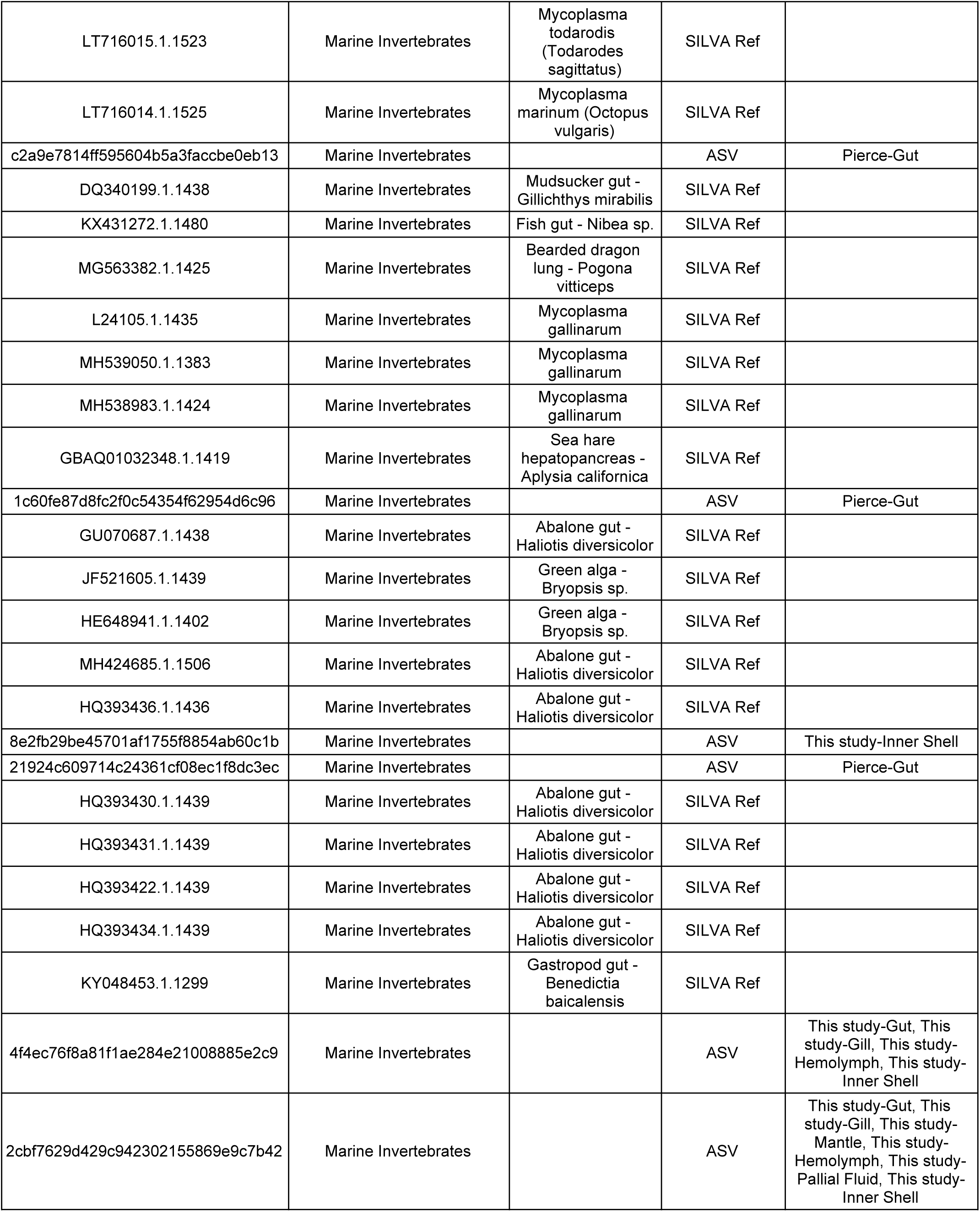
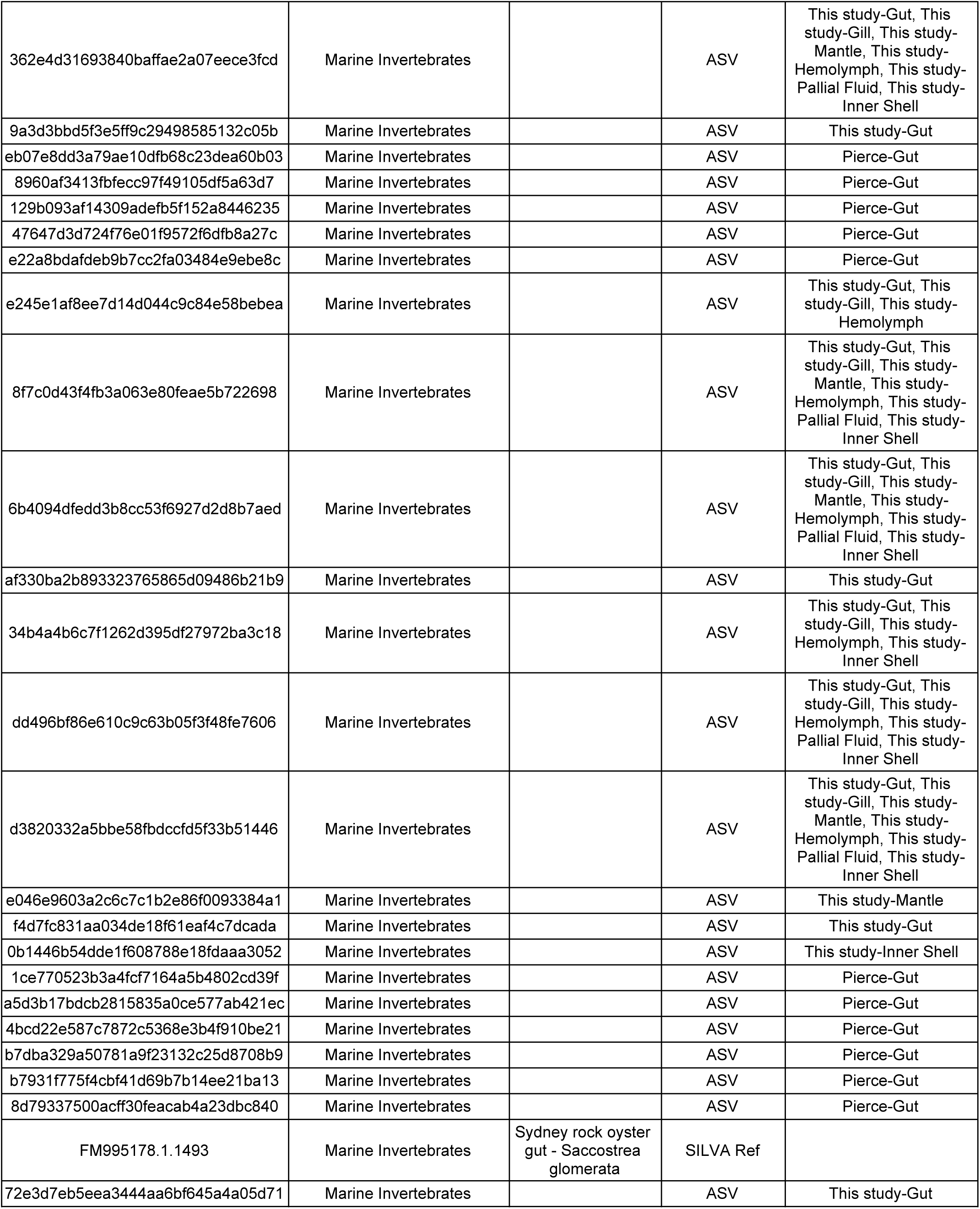
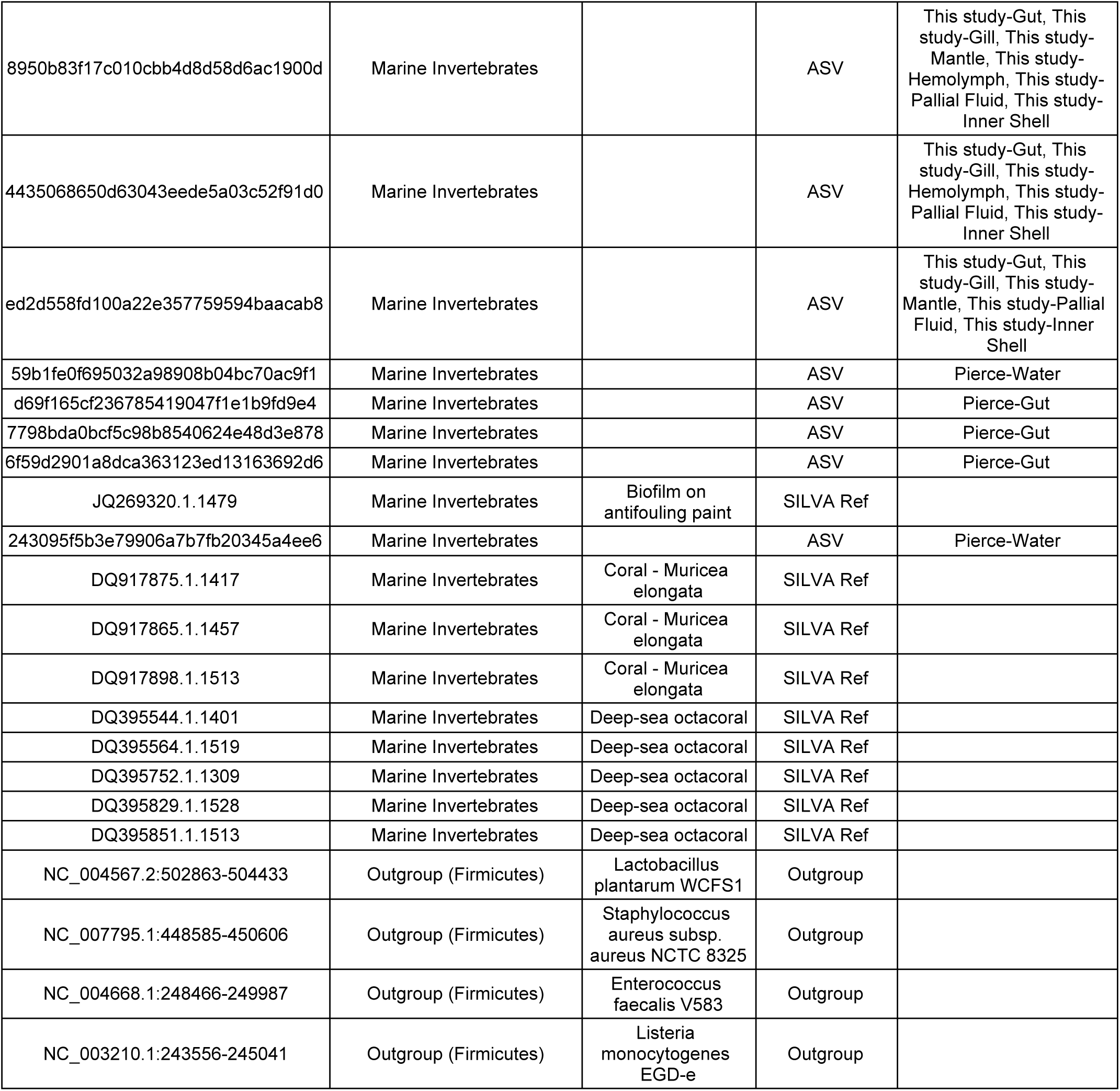
Metadata showing the references and ASVs included in the phylogeny of **Figure 4**. Reference genes were labeled as identifiers in the SILVA reference sequence database. ASVs were labeled as their unique identifiers. The Clade assignments indicate the positioning of a reference or ASV to a specific clade in **Figure 4**. More specific information about the source environment of each reference gene was represented in the Environment column. The source samples (this study vs. [39]) for each ASV were indicated in the Sample Types column.

**Table S8.**
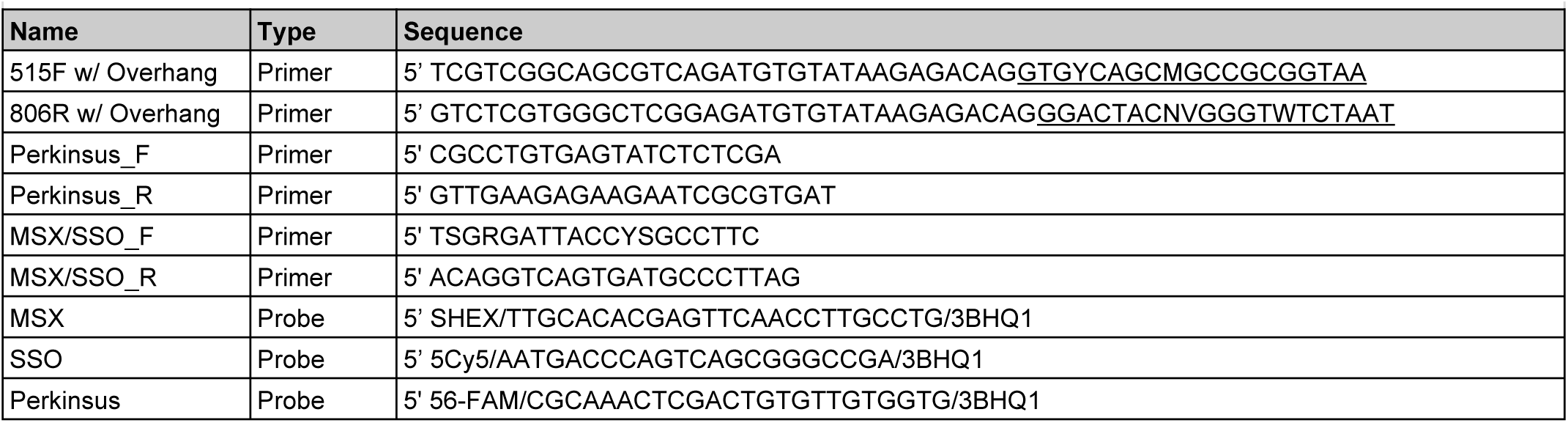
Summary of primers and probes used in this study. The locus-specific part of the primers 515F w/ Overhang and 806R w/ Overhang are underlined while the rest of the sequence contains a sequencing adapter.

**Table S9.**
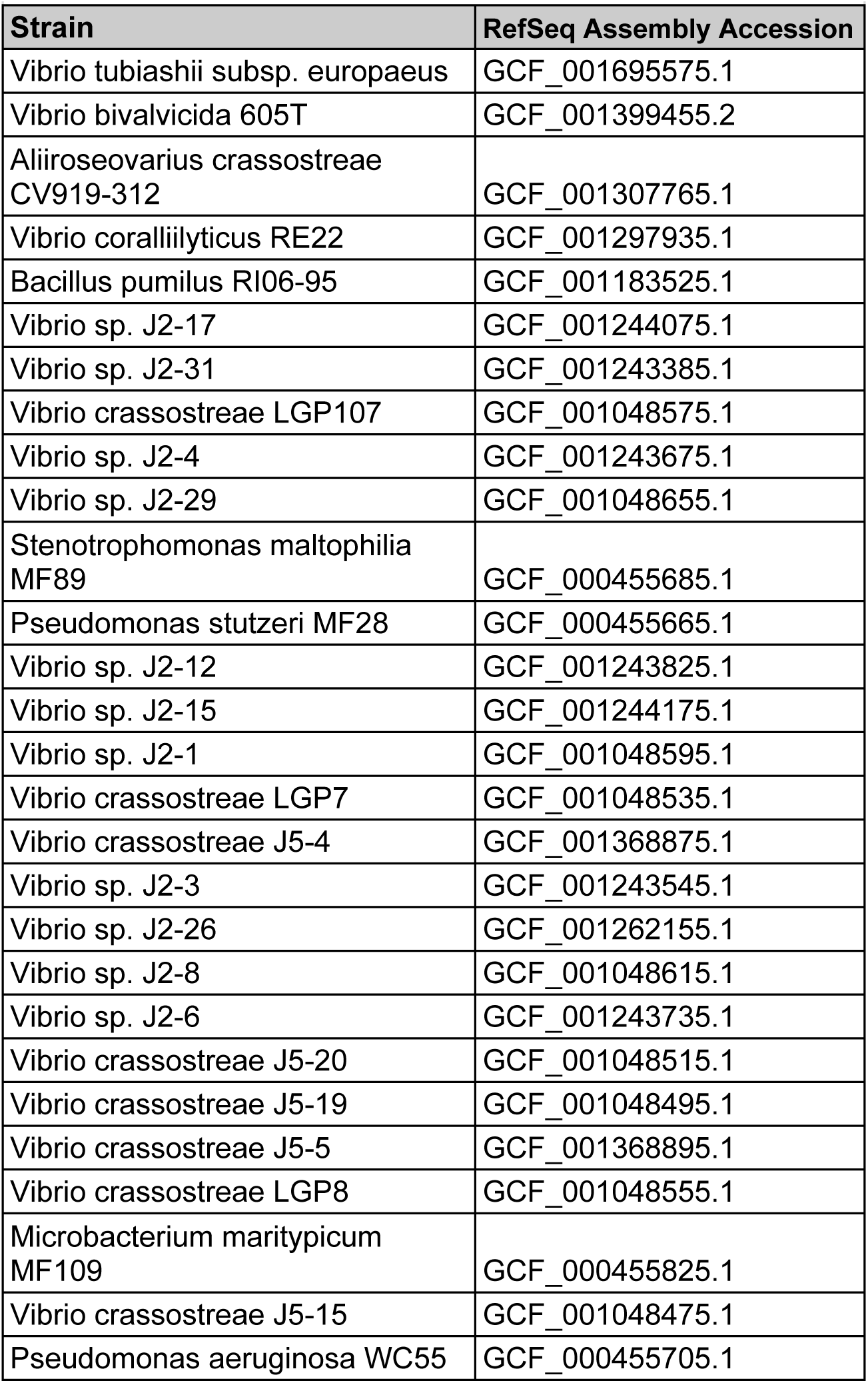
Strain names and NCBI RefSeq assembly accessions for a set of manually curated draft genomes of oyster derived isolates.

